# The non-canonical thioreductase TMX2 is essential for neuronal survival during embryonic brain development

**DOI:** 10.1101/2024.06.18.599494

**Authors:** Jordy Dekker, Wendy Lam, Herma C. van der Linde, Floris Ophorst, Charlotte de Konink, Rachel Schot, Gert-Jan Kremers, Leslie E. Sanderson, Woutje M. Berdowski, Geeske M. van Woerden, Grazia M.S. Mancini, Tjakko J. van Ham

## Abstract

Biallelic variants in thioredoxin-related transmembrane 2 protein (TMX2) can cause a brain malformation of cortical development (MCD), characterized by primary microcephaly, polymicrogyria and pachygyria by an unknown mechanism. To better understand and visualize how TMX2 loss disrupts brain development *in vivo* we investigated the function of TMX2, using the zebrafish embryo as a model system. We generated zebrafish deficient for *TMX2* ortholog *tmx2b*, which during the first 2 days post fertilization (dpf) showed normal behavioral activity and brain developmental hallmarks. From 3 dpf onwards however, *tmx2b* mutants failed to exhibit locomotor activity, which was accompanied by cell death primarily in the brain, but not in other organs or in the spinal cord. Strikingly, cell death in *tmx2b* mutants occurs specifically in newborn neurons within a ∼1.5-hour timeframe, whereas neuronal progenitor and other glial cells are preserved, and could be suppressed by inhibiting neuronal activity. *In vivo* GCaMP6s calcium imaging showed a persistent ∼2-fold increase in calcium in neurons after the onset of cell death. This suggests that calcium homeostasis underlies the *tmx2b* mutant brain phenotype. Altogether, our results indicate that TMX2 is an evolutionary conserved, protective regulator essential specifically for newborn neurons to survive after their differentiation in the vertebrate embryonic brain.

## Introduction

Embryonic development of the human cerebral cortex is an intricate process directed by an interplay of molecular genetic and environmental factors. Disruption of these factors impacts normal cortical development and can result in a malformation of cortical development (MCD) with a neurodevelopmental disorder (NDD), often contributing to a global developmental delay and epilepsy with a high disease burden for the affected individual and their families (1, 2). The last 30 years many genetic causes for MCD have been identified and functional studies reveal that disruption of diverse molecular pathways—such as cell division, ER stress and the cytoskeletal regulation—can impair cortical development (1, 3-5). This genetic heterogeneity has not only advanced our understanding of MCD development, but also significantly expanded our knowledge on the molecular pathways involved in normal cortical development of the human (1, 5).

*TMX2*, coding for thioredoxin (TRX)-related transmembrane 2 (*TMX2*) protein, is ubiquitously expressed in humans; however, biallelic variants in *TMX2* have been associated with severe NDD including epileptic encephalopathy, microcephaly, and cortical malformations such as unlayered polymicrogyria and pachygyria (6-8). TMX2 is a member of the protein disulfide isomerase (PDI) family, consisting over 20 endoplasmic reticulum (ER) chaperones that assist in protein folding by alternating intra- and inter-molecular cysteine residues between oxidized and reduced states (8-14). TMX2 is one of five membrane-tethered PDIs, along with TMX1, TMX3, TMX4 and TXNDC15, collectively comprising the thioredoxin-related transmembrane (TMX) protein subfamily within the PDI family (8, 13). The TMX proteins share a N-terminal signal peptide required for ER targeting, a single transmembrane domain, a TRX domain containing the active site with two cysteine residues (CXXC) and a C-terminal ER retention motif (13). Unlike the other TMX proteins, TMX2 consists of multiple transmembrane domains and harbors an atypical active site, with the N-terminal cysteine being replaced by a serine residue (SNDC, instead of CXXC), and a TRX domain that is directed towards the cytosol (6, 13, 15). As oxidoreductase activity requires a canonical CXXC motif, containing two cysteine residues, it is uncertain whether TMX2 exerts such a function (10, 13, 16).

Although oxidoreductase activity of TMX2 lacks definitive proof, its interaction with ER chaperones and proteins of the unfolded protein response (UPR) hint towards a putative role in protein folding by TMX2 (6). *In vitro* studies in mouse cortical neurons and human cholangiocarcinoma cells (HuCCT1) showed that *TMX2* knockdown affects expression levels of UPR proteins (17, 18). Furthermore, TMX2 was found to localize to mitochondria-ER contacts (MERCs). Interactome analysis of TMX2 in human HEK293T cells showed several MERC located Ca^2+^ chaperons and channel proteins (e.g. calnexin, RCN2, SERCA2) as main interactors, suggesting that TMX2 could potentially also modulate ER-mito-chondria Ca^2+^-regulated crosstalk, similar to TMX1 and other PDIs (6, 19-21). Cultured skin fibroblasts of individuals with pathogenic variants in *TMX2* exhibited mitochondrial dysfunction, further stressing an important role for TMX2 in mitochondrial physiology (6). TMX2 also functions at the nuclear pore where it regulates nuclear transport via importin-β (15). Disturbance of these processes have been implicated in disrupting cortical development, hence emphasizing that the physiological function of TMX2 is essential for normal brain development (4, 22-24)

Currently, it is unknown how TMX2 loss affects cortical development *in vivo* and causes microcephaly and cortical malformation, nor how TMX2 normally regulates human brain development. The use of animal embryos has been instrumental in recapitulating steps of human brain development and particularly zebrafish embryo has successfully been explored to study the effects of human pathogenic variants (25, 26). To obtain a better understanding we generated zebrafish deficient for the *TMX2* ortholog, *tmx2b*. The *tmx2b* deficient zebrafish show normal embryonic developmental hallmarks in the first two days post fertilization (dpf). At 3 dpf *tmx2b* KO zebrafish show a rapid onset massive neuronal cell death affecting most of the brain, which appears to not progress further by 5 dpf. Cell death is restricted to post-mitotic neurons as glia cell populations are unaffected. Calcium imaging suggested a possible dysregulation of Ca^2+^ in the neurons of *tmx2b* KO zebrafish. Our results indicate that TMX2 has a protective role in neurons by regulating Ca^2+^ concentrations in the brain.

## Material and methods

### Zebrafish housing and husbandry

Zebrafish were under standard housing and husbandry conditions (27). The adult animals were fed twice a day on a 14h-10h light-dark cycle. Zebrafish embryos and larvae were kept at 28 °C in E3 medium buffered with 20 mM HEPES (pH 7.2) (referred to as E3) on a 14h-10h light-dark cycle till 5 days post fertilization (dpf). To prevent pigmentation E3 medium was changed to 0.003% (m/v) 1-phenyl 2-thiourea (PTU; Sigma-Aldrich) at 24 hours post fertilization (hpf). 50 zebrafish embryos/larvae were kept in a petri dish containing 25 mL E3 medium. The transgenic zebrafish lines used in this study are listed in **Table S1** (28-35). The experiments at a defined dpf in this manuscript were always performed in the afternoon between 14:00 and 17:00. So an experiment performed at a 3 dpf zebrafish larva was performed at an embryo 72-75 hpf of age.

### CRISPR-Cas9 genome editing

Alt-R® CRISPR-Cas9 crRNAs were designed and ordered via integrated DNA technologies (IDT) website. The crRNA sequence targeted against exon 1 of *tmx2a* is 5’-GGAGTCTCCGTCCTCTCTCT-3’ and exon 3 of *tmx2b* 5’-CGTTGGCCACTTTACAGAAG-3’’. Equal volumes of Alt-R® CRISPR-Cas9 tracrRNA and crRNA in duplex buffer were incubated at 95°C for 5 minutes, followed by cooling down at room temperature (RT) to allow cr:tracrRNA complex formation. The Sp-Cas9 plasmid (Addgene plasmid #62731) was utilized to synthesize SpCas9 as described previously (36). For the formation of cr:tracrRNA-Cas9 ribonucleoproteins, 25 pmol SpCas9 was mixed with 50 pmol cr:tracrRNA and incubated at RT for 5 minutes. 3 uL 300 mM KCL and 0.3 uL phenol red were subsequently added, and 1 nl was injected in the 1-cell stage of WT zebrafish embryo. Indel frequency was determined by Sanger sequencing of genomic DNA extracted from tail fins of adult zebrafish and analysis of sequencing files in TIDE (37)Cultured</keyword><keyword>DNA Mutational Analysis/*methods</keyword><keyword>Genomics/methods</keyword><keyword>Humans</keyword><keyword>*INDEL Mutation</keyword><keyword>K562 Cells</keyword><keyword>Mutagenesis</keyword><keyword>Polymerase Chain Reaction</keyword><keyword>Software</keyword></keywords><dates><year>2014</year><pub-dates><date>Dec 16</date></pub-dates></dates><isbn>1362-4962 (Electronic. Primers used are listed in **Table S2**. Founder zebrafish positive for indels in *tmx2a* and *tmx2b* respectively were outcrossed against WT zebrafish to generate a heterozygous F1 generation. Genotypes of F1 adult zebrafish were determined by Sanger sequencing. Zebrafish heterozygous for a 1 bp insertion in *tmx2a* and a 7 bp insertion in *tmx2b* were selected.

### Genotyping zebrafish embryo *tmx2b* by allele specific PCR

Since homozygous loss of *tmx2b* was embryonically lethal we performed all experiments on an incross of heterozygous (*tmx2b*^*+/-*^) zebrafish. The WT and heterozygous zebrafish siblings (*tmx2b*^*+/?*^) served as the control group against the *tmx2b*^*-/-*^. After each experiment zebrafish embryo and larvae were euthanized and DNA extraction was performed by lysis of embryo in 40 uL 50 mM NaOH, followed by incubation at 95°C for 30 minutes. Samples were cooled down to RT and 4 uL 1 M Tris-HCl pH8.0 was added. Allele specific PCR consisted of three different primers, a forward and reverse primer with in the middle an allele specific primer able to bind either only the WT or mutant allele **(Table S2)**. Allele specific PCR with FastStart Taq (Roche diagnostics) was performed under following conditions: 95 °C for 5 min, 10 cycles of 94 °C for 30 s, touchdown 65 → 60 °C (-0.5 °C/cycle) for 30 s, 72 °C for 45 s, followed by 25 cycles of 94 °C for 30 s, 60 °C 30 s, 72 °C for 45 s and finished by a final step of 72 °C for 5 min. 3 uL PCR product was visualized on 2% agarose gel in Tris-acetate-EDTA buffer.

### Touch response analysis

Zebrafish embryos were placed individually into 48-Wells plate containing 1 mL E3 medium. Touch response was provoked with a plastic loading pipette tip at 1 dpf. At 2 dpf zebrafish embryo were first dechorionated and touch response was provoked with a 23G needle from 2 till 4 dpf. Touch responses were provoked a maximum of three times per zebrafish and recorded as, 1) normal: upon first touch zebrafish swims out of view, 2) delayed: >1 provocations before effective swimming or ineffective swimming response as observed by only twitching of the body, or 3) no response: after 3 provocations no movement visible. Videos were recorded with the Olympus SZX116 microscope (Olympus) with a DP72 camera (Olympus).

### Neutral Red staining

Zebrafish larvae at 3 and 5 dpf were incubated in 2.5 ug/mL Neutral Red (Sigma-Aldrich) dissolved in E3 containing 0.003% (m/v) PTU for 2h at 28°C. After incubation zebrafish were washed and incubated in E3 containing 0.003% (m/v) PTU for 20 minutes at 28°C. Stained zebrafish larvae were anesthetized with 0.016% (m/v) Ethyl 3-aminobenzoate methanesulfonate salt (Tricaine; Sigma-Aldrich, A5040) and embedded in 1.8% low melting point agarose (Invitrogen) and subjected to imaging.

### Lysotracker staining

Two times fifteen zebrafish larvae at 2,3 and 5 dpf were transferred to a round bottom 2 mL Eppendorf tube. 1 mM LysoTracker™ Red DND-99 (Invitrogen, L7528) was diluted 1:100 with E3 containing 0.003% (m/v) PTU. Subsequently, E3 medium was removed from the round bottom 2 mL Eppendorf tube and zebrafish were incubated in 250 uL of the 10 **μ**M Lysotracker solution at 28°C for 40 minutes in the dark with the caps opened. Next, zebrafish were washed and incubated in E3 containing 0.003% (m/v) PTU for 20 minutes at 28°C in the dark, before embedding and imaging.

### Drug treatments

Zebrafish eggs were collected shortly after the first eggs were laid, within a 15-minute time window. Details on drugs, start of treatment, concentrations and drug refreshments are listed in **Table S3**. During drug treatment 50 zebrafish embryo/larvae were kept in a total volume of 25 mL to assure normal embryonic development. At 3 dpf drugs were removed and zebrafish were subjected to imaging.

### Image acquisition

For body length and brightfield images of lateral view head the zebrafish were first anesthetized in 0.016% Tricaine and transferred to 10% methylcellulose (Sigma, M7027), allowing correct positioning of the zebrafish during imaging. Images of total zebrafish and lateral view head were acquired with Leica M165 FC microscope using a 10x dry objective and a Leica DFC550 camera. For neutral red images the same microscope was used and serial images (2-4 images per zebrafish brain) in the z-plane were acquired.

*in vivo* confocal imaging of zebrafish was performed at a Leica SP5 intravital microscope containing a 20x/1.0 NA water dipping objective using 488 (GFP), 514 (mVenus) and 561 (DsRED, Lysotracker) lasers. Z-stacks with z-step size ranging from 1-4 **μm** were acquired.

### Time-lapse imaging

For overnight time-lapse imaging of the excitatory (*vglut2:*DsRED) and inhibitory neurons (*gad1b:*GFP) and Ca^2+^ imaging (*elavl3:*NLS-GCaMP6s) 55 hpf zebrafish embryo were anesthetized with 0,5 mg/ml α-bungarotoxin (Invitrogen, B1601). This toxin binds irreversibly to the neuromuscular junctions (nicotinic acetylcholine receptors); thereby, persevering normal (brain) development of the zebrafish embryo. Zebrafish were embedded in 0.8% low melting point agarose in the InViSPIM lattice pro sample holder. Overnight imaging was performed with the InViSPIM lattice pro (Bruker) at 28 °C. For both excitatory and inhibitory neurons every 10 minutes the entire brain of each zebrafish was imaged through the z-plane from 63 hpf till 75 hpf.

### Image and statistical analysis

All images were processed and analyzed before genotyping with Fiji ImageJ software. Total body lengths were measured from jaw to fin tail in zebrafish imaged from dorsal side. Apoptotic clusters (*ubb:*SecA5-mVenus+ clusters) and oligodendrocyte precursor cells (OPCs; *olig1:*NLS-mApple) were analyzed with the 3D object counter plugin. Threshold were same for each image of each experiment and dpf, voxel size minimum value was 4. For excitatory (*vglut2:*DsRED) and inhibitory neurons (*gad1b:*GFP) a threshold was set on a z-projection of the brain. The same threshold was used for each image of each experiment and dpf, and total midbrain area was calculated. Excitatory and inhibitory neurons numbers were manually counted in a region of interest in the middle of the spinal cord images. Radial glial cell fibers (*her4*.*3:*EGFP) were manually counted in a region of interest in the right hemisphere of the midbrain. Microglia (Neutral red+ or *mpeg1:*EGFP+) were manually counted in the whole midbrain area. Lysotracker and Ca^2+^ (*elavl3:*NLS-GCaMP6s) intensities were measured as the mean fluorescence intensity on Z-projections of the midbrain area. To analyze Lysotracker area in microglia and microglia (*mpeg1:*EGFP+) circularity a threshold was applied on the Z-projections of midbrain regions. Six microglia per zebrafish brain were selected for morphological analysis and Lysotracker area calculations.

## Results

### Generation of *tmx2a* and *tmx2b* knockout zebrafish

To investigate the impact of TMX2 loss *in vivo* we generated a genetic zebrafish model by mutating both *TMX2* homologs, *tmx2a* and *tmx2b*, referred to as: *tmx2a*^*-/-*^ and *tmx2b*^*-/-*^, by introducing frameshifting mutations in exon 1 and 3 respectively **(Figure S1)**. If both these mutated transcripts are not degraded by nonsense mediated RNA decay, translation would result in a truncated protein lacking the complete catalytic thioredoxin-like (TRX) domain, preserving only the signal peptide and first transmembrane domain **(Figure S2A)**. We observed by visual inspection that adult *tmx2a*^*-/-*^ zebrafish did not differ from controls in growth and survival, while *tmx2b*^*-/-*^ zebrafish could only be kept in heterozygous state due to early lethality after reaching the feeding stage (data not shown). Tmx2a and Tmx2b share respectively 64% and 70% protein homology with TMX2 **(Figure S2B**,**C)**. Tmx2a and Tmx2b share 67% homology and a distance tree (BLOSUM62 algorithm) showed that Tmx2b is phylogenetically closer related to TMX2 than Tmx2a **(Figure S2C)**. Furthermore, in control zebrafish brain at 5 days post fertilization (dpf) only *tmx2b* is expressed and *tmx2a* is not (Average transcripts per million (TPM): *tmx2b*: >53.5; *tmx2a*<0.001) and the zebrafish RNA-seq database (zfRegeneration) showed that across multiple tissues and embryonic/ larval developmental stages *tmx2b* is the main expressed gene **(Figure S3A**,**B)** (38). Altogether, these data show that Tmx2b is most equivalent to TMX2 and most relevant to its function in the brain; therefore, we decided to use *tmx2b*^*-/-*^ zebrafish to model TMX2 deficiency **(Figure 1A,B)**.

**Figure 1.**
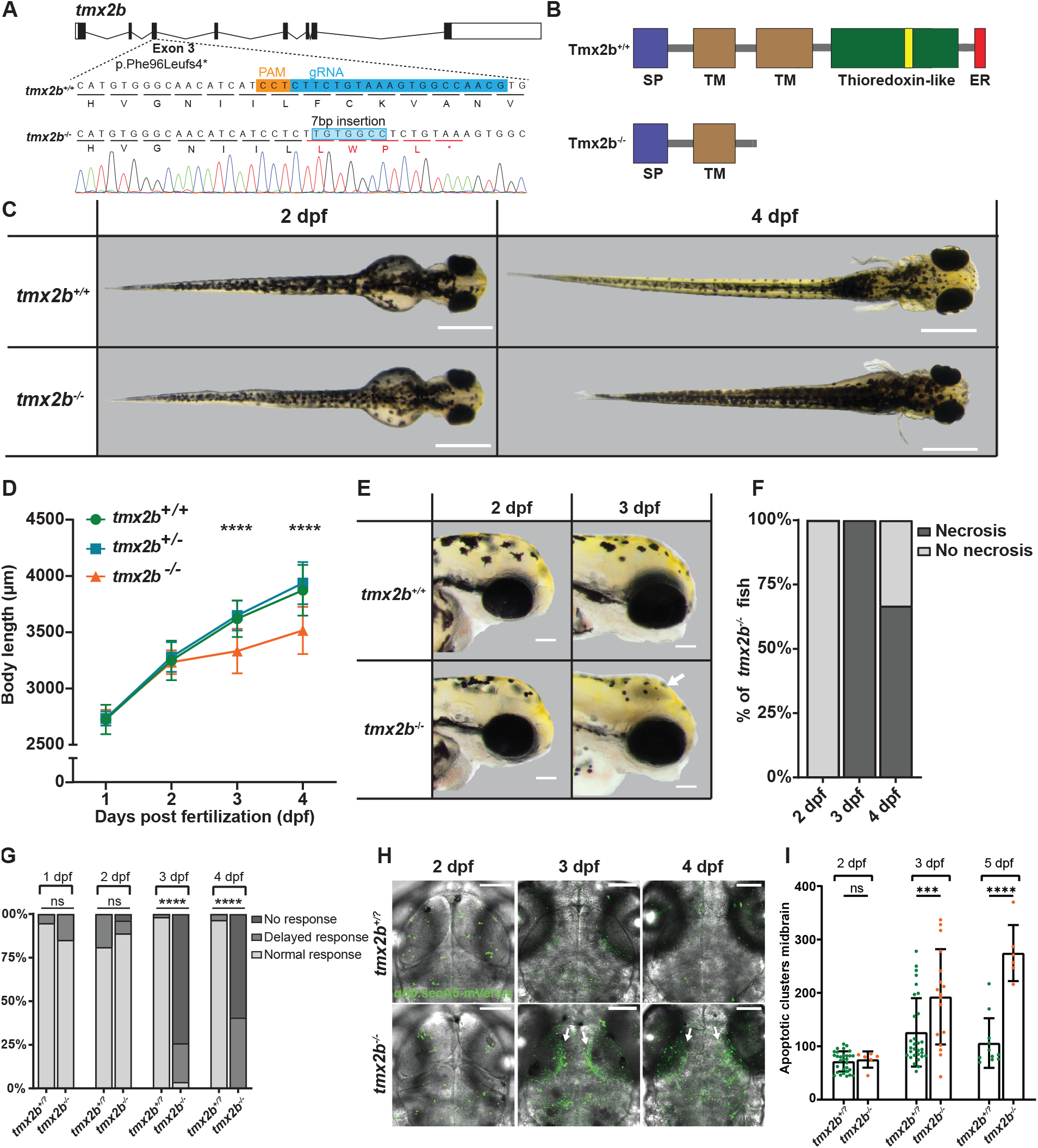
*txm2b*^*-/-*^ zebrafish display a developmental regression from 3 dpf with cell death in brain regions. **(A)** Schematic representation of the *tmx2b* gene (Ensembl transcript ID:ENSDART00000009858.6) with the gRNA target in exon 3. Sanger sequence data of *tmx2b*^*-/-*^ zebrafish embryo with a homozygous c.285_286insTGTGGCC, p.Phe96Leufs4* mutation. **(B)** Schematic representation of the Tmx2b protein. *tmx2b*^*-/-*^ zebrafish lack the thiore-doxin-like domain containing the catalytic S-X-X-C motif (yellow box) of Tmx2b. SP, signal peptide sequence; TM, transmembrane domain; ER, ER-retention motif. **(C)** Representative images of same *tmx2b*^*+/+*^, *tmx2b*^*+/-*^ and *tmx2ab*^*-/-*^ zebrafish at 1 to 4 days post fertilization (dpf). *tmx2b*^-/-^ have normal body morphology till 2 dpf and from 3 dpf onwards display a developmental decline. Scale bars indicate 500 μm. **(D)** Body length measurements of *tmx2b*^*+/+*^, *tmx2b*^*+/-*^ and *tmx2b*^*-/-*^ zebrafish from 1-4 dpf. Data are represented as mean ± SD. N=2 experiments, *tmx2b*^*+/+*^, n=9; *tmx2b*^*+/-*^, n=37; *tmx2b*^*-/-*^, n=15. Two-way ANOVA with Tukey’s multiple comparisons test. **(E)** Representative images of *tmx2b*^*+/+*^, *tmx2b*^*+/-*^ and *tmx2b*^*-/-*^ zebrafish brightfield images lateral view of head. *tmx2b*^*-/-*^ zebrafish develop a gray discoloration in brain region (white arrow) indicative of necrosis. This gray discoloration is not always present at 4 dpf in *tmx2b*^*-/-*^ zebrafish. Scale bars indicate 100 μm. **(F)** Quantification of same *tmx2b*^*-/-*^ zebrafish with gray discoloration (necrosis) at 2 till 4 dpf. At 3 dpf all zebrafish *tmx2b*^*-/-*^ all zebrafish have necrosis. At 4 dpf 33% of the *tmx2b*^*-/-*^ zebrafish no longer have visible necrosis in the brain. N=2 experiments, n=15 zebrafish. **(G)** Quantification of *tmx2b*^*+/?*^ and *tmx2b*^*-/-*^ zebrafish with a normal/delayed or absent touch response from 1 till 4 dpf. N=2 experi-ments, *tmx2b*^*+/?*^ n=58, *tmx2b*^*-/-*^ n=27 zebrafish. Fisher’s exact test (delayed and no touch response groups were combined for statistical test). **(H)** Representative images of *ubb:*secA5-mVenus+ (green) apoptotic clusters merged with brightfield image of *tmx2b*^*+/?*^ and *tmx2b*^*-/-*^ zebrafish at 2, 3 and 4 dpf. Increased apoptosis is mostly pronounced in optic tecti (white arrows). Scale bars indicate 100 μm. **(I)** Quantification of apoptotic clusters brain of *tmx2b*^*+/?*^ and *tmx2b*^*-/-*^ zebrafish at 2, 3 and 4 dpf. Data are represented as mean ± SD. 2,3,4 dpf: N=2,2,1 experiments; tmx2b^+/?^, n=30,30,11; tmx2b^-/-^, n=6,9,6. Two-way ANOVA, Šídák›s multiple comparison test. ***p < 0.001, ****p < 0.0001.

### *tmx2b*^-/-^ zebrafish embryo exhibit developmental regression from 3 days post fertilization

As *tmx2b*^*-/-*^ zebrafish did not reach adulthood and never survived beyond 5-10 dpf, we analyzed total body length at embryonic stages from 1-4 dpf as a marker of general development (39, 40). At 1 and 2 dpf *tmx2b*^-/-^ zebrafish embryo appeared normal and body length did not differ from controls **(Figure 1C,D and Figure S4A,B,C)**. At 3 and 4 dpf *tmx2b*^*-/-*^ zebrafish displayed developmental decline and were shorter than *tmx2b*^*+/+*^ and *txm2b*^*+/-*^ zebrafish **(Figure 1C,D and Figure S4A,D,E)**. Additionally, *tmx2b*^*-/-*^ zebrafish at 3 and 4 dpf had a gray discoloration in the brain, a sign of necrotic cell death **(Figure 1E and Figure S4F)** (41). Interestingly, at 4 dpf ∼33% of the *tmx2b*^*-/-*^ zebrafish seemed to have resolved the visible necrosis, suggesting a transient phase of cell death **(Figure 1F)**. Heterozygous *tmx2b*^*+/-*^ zebrafish had normal body length, did not exhibit brain necrosis and showed no visually discernable phenotype; therefore, we combined *tmx2b*^*+/+*^ and *tmx2b*^*+/-*^ as one control group (referred to as *tmx2b*^*+/?*^) for the subsequent experiments **(Figure 1C-F)**. At 1 and 2 dpf *tmx2b*^*-/-*^ mutants responded normally to touch, but were non-responsive at 3 and 4 dpf, indicating that disease onset in *tmx2b*^*-/-*^ appeared sudden between 2 and 3 dpf. **(Figure 1G and Figure S5 and Videos S1-9)**. The first *tmx2b*^*-/-*^ zebrafish started to develop the gray discoloration when the *tmx2b*^+/?^ zebrafish siblings were between the pec-fin (60 hpf) and protruding-mouth stage (72 hpf), indicating that brain necrosis initiates at the end of the embryonic phase **(Figure S4G)**.

Next, we assessed apoptosis and visualized apoptotic cells (*ubb:*SecA5-mVenus+ clusters) in zebrafish brain at 2, 3 and 4 dpf **(Figure 1H,I and Figure S6)**. At 2 dpf numbers of apoptotic clusters in *tmx2b-/* and *tmx2b*^*+/?*^ zebrafish did not differ **(Figure 1H,I and Figure S6)**. Consistent with previous findings we observed increased numbers of apoptotic clusters in *tmx2b-/* compared to *tmx2b*^*+/?*^ zebrafish at 3 and 4 dpf **(Figure 1H,I and Figure S6)**. These data indicate that loss of Tmx2b does not affect general development the first 2 dpf, whereas at 3 dpf *tmx2b*^*-/-*^ zebrafish all develop growth regression and massive cell death in the brain that appears to be non-progressive up to 4 dpf and their inability to voluntarily move after 3 dpf

### Tmx2b loss causes neuronal cell death at 3 dpf in zebrafish embryo brain

Since we observed the massive cell death in the brain we focused on which brain cell types were affected by Tmx2 loss. Since, many individuals with biallelic pathogenic LoF variants in *TMX2* develop a primary microcephaly (i.e. present at birth) with abnormal cortex structure (polymicrogyria) and some show progressive microcephaly also after birth, we hypothesized that both the radial glial cells (neural and glial progenitor cell) and the generated neurons undergo cell death in *tmx2b*^*-/-*^ (1, 2, 6, 7). First, we determined the effect of Tmx2b loss in the excitatory (*vglut2:*DsRED) and inhibitory neurons (*gad1b:*GFP). At 2 dpf the excitatory and inhibitory neurons were similar between *tmx2b*^*-/-*^ and *tmx2b*^*+/?*^ **(Figure 2A-D and Figure S7&8)**. At 3 dpf both excitatory and inhibitory neuronal cell loss was observed within the midbrain in *txm2b*^*-/-*^ larvae **(Figure 2A-D and Figure S7&8)**. Brain size did not differ at 3 dpf between *tmx2b*^*-/-*^ and *tmx2b*^*+/?*^ zebrafish, indicating that the excitatory and inhibitory neurons were present in the brain at 2 dpf and died between 2 and 3 dpf **(Figures S7&S8)**. Neuronal cell death was not progressive as the area positive for excitatory neurons increased between 3 and 5 dpf in *tmx2b*^*-/-*^ zebrafish; however, the loss of inhibitory neurons appeared progressive between 3 and 5 dpf **(Figure 2A-D and Figure S7&8)**. As Tmx2 deficiency is early lethal in zebrafish and embryonic lethal in mice, we tested whether Tmx2 is important for survival of mammalian neurons *in vitro*.

**Figure 2.**
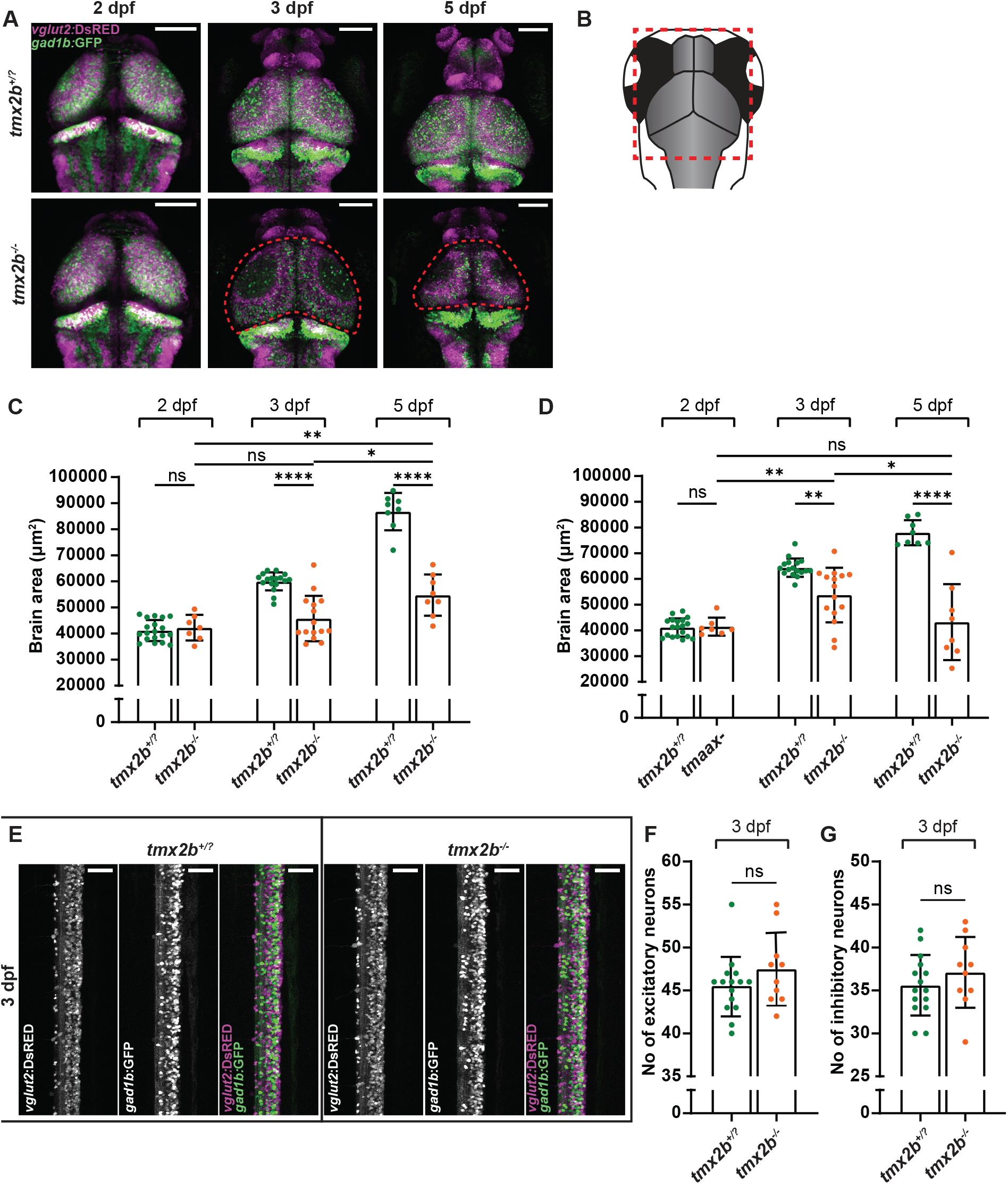
Neurons in central brain regions *tmx2b*^*-/-*^ zebrafish undergo cell death at 3 dpf. **(A)** Representative images of *vglut2:*DsRED^+^ excitatory neurons (magenta) and *gad1b:*GFP^+^ inhibitory neurons (green) in *tmx2b*^*+/?*^ and *tmx2b*^*-/-*^ zebrafish brain at 2, 3 and 5 dpf. Red boxes indicate areas of neuronal cell loss. **(B)** Quantification of excitatory neuron brain area in *tmx2b*^*+/?*^ and *tmx2b*^*-/-*^ zebrafish at 2, 3 and 5 dpf. Excitatory neuronal cell death is observed at 3 dpf in *tmx2b*^*-/-*^ zebrafish, which shows a minor recovery at 5 dpf. **(C)** Quantification of inhibitory neuron brain area in *tmx2b*^*+/?*^ and *tmx2b*^*-/-*^ zebrafish at 2, 3 and 5 dpf. Contrary to the excitatory neurons, inhibitory neuronal loss is progressive from 3 dpf onwards. **(A-C)** 2,3,5 dpf: N=1,2,1 experiments; *tmx2b*^*+/?*^, n=19,17,8; *tmx2b*^*-/-*^, n=7,15,8. Scale bars indicate 100 μm. Two-way ANOVA, Tukey’s multiple comparisons test. **(F)** Representative images of *vglut2*:DsRED+ excitatory neurons and *gad1b:*GFP+ inhibitory neurons in spinal cord of *tmx2b*^*+/?*^ and *tmx2b*^*-/-*^ zebrafish at 3 dpf. **(G)** Quantification of numbers of excitatory neurons in spinal cord of in *tmx2b*^*+/?*^ and *tmx2b*^*-/-*^ zebrafish at 3 dpf. **(H)** Quantification of numbers of inhibitory neurons in spinal cord of in *tmx2b*^*+/?*^ and *tmx2b*^*-/-*^ zebrafish at 3 dpf **(F-H)** N=2 experiments, *tmx2b*^*+/?*^ n=15, *tmx2b*^*-/-*^ n=10 zebrafish. Scale bars indicate 50 μm. One-way ANOVA with Šídák›s multiple comparisons test. Data are represented as mean ± SD. *p < 0.05, **p < 0.01, ***p < 0.001, ****p < 0.0001.

Since the abnormal brain discoloration was only observed in the 3 dpf brain we assessed whether neuronal loss also occurred in the neurons of the spinal cord. Both excitatory and inhibitory cells numbers in *tmx2b*^*-/-*^ were comparable to *tmx2b*^*+/-*^ larvae **(Figure 2F-H and Figure S9)**. These data indicate that Tmx2b loss causes both excitatory and inhibitory cell death between 2 and 3 dpf limited to the brain regions, as the neurons in the spinal cord are unaffected.

### Glia cell populations are not directly affected by Tmx2 loss

Next, we determined whether cell death was restricted to neuronal populations or whether glial cell populations were also affected. First, we assessed the number of radial glial cells (*her4*.*3:*EGFP), which exert both astrocytic functions and neural stem cell properties in zebrafish (42, 43). Surprisingly, the radial glial cell numbers in *tmx2b-/* did not differ from *tmx2b*^*+/?*^ control zebrafish at 2, 3 and 5 dpf **(Figure 3A,B)**. Although radial glial cell numbers were similar, we did observe that the soma of radial glial cells, which normally resides at the apical edge of the ventricular zone, migrated from the ventricular zone towards the midbrain/optic tectum **(Figure 3A)**.

**Figure 3.**
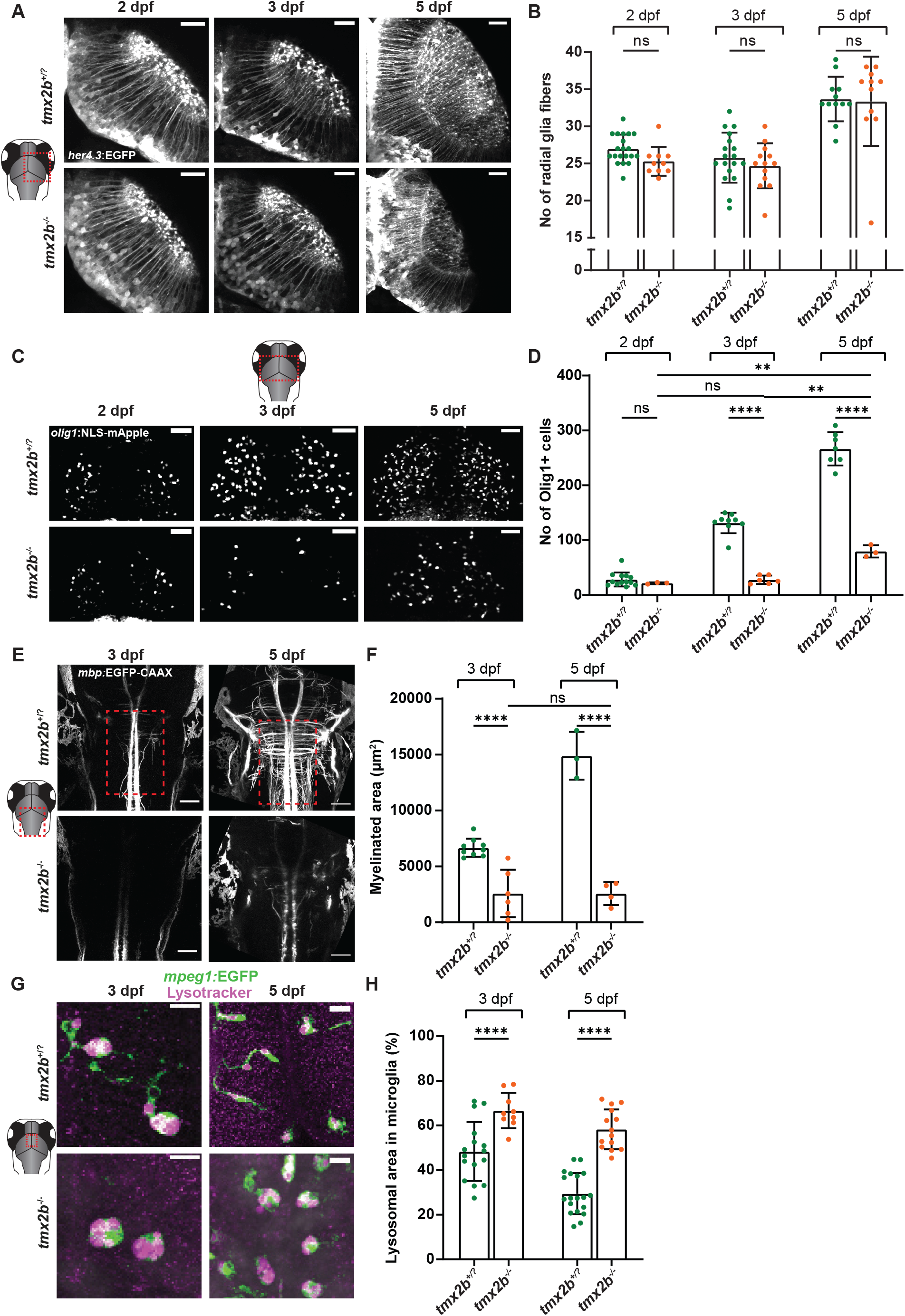
Glia cell populations in *tmx2b*^*-/-*^ zebrafish. **(A)** Representative images of *her4*.*3:*EGFP+ radial glia cells in right midbrain hemisphere of *tmx2b*^*+/?*^ and *tmx2b*^*-/-*^ zebrafish at 2, 3 and 5 dpf. Scale bars indicate 25 μm. **(B)** Quantification of number of radial glial fibers right midbrain hemisphere of *tmx2b*^*+/?*^ and *tmx2b*^*-/-*^ zebrafish at 2, 3 and 5 dpf. 2,3,5 dpf: N=1,1,1 experiments; *tmx2b*^*+/?*^, n=20,18,12; *tmx2b*^*-/-*^, n=11,13,11. Two-way ANOVA, Šídák›s multiple comparisons test. **(C)** Representative images of *olig1:*NLS-mApple (magenta) OPCs in the midbrain of *tmx2b*^*+/?*^ and *tmx2b*^*-/-*^ zebrafish at 2, 3 and 5 dpf. Scale bars indicate 100 μm. **(D)** Quantification of OPCs in midbrain of *tmx2b*^*+/?*^ and *tmx2b*^*-/-*^ zebrafish at 2, 3 and 5 dpf. 2,3,5 dpf: N=1,1,1 experiments; tmx2b^+/?^, n=13,9,7; tmx2b^-/-^, n=3,6,3. Two-way ANOVA, Tukey’s multiple comparisons test. **(E)** Representative images of *mbp:*EGFP-CAAX in hindbrain region of *tmx2b*^*+/?*^ and *tmx2b*^*-/-*^ zebrafish at 3 and 5 dpf. Scale bars indicate 50 μm. **(F)** Quantification of myelinated area hindbrain of *tmx2b*^*+/?*^ and *tmx2b*^*-/-*^ zebrafish at 3 and 5 dpf. Measured area is indicated by red dashed box in **(E). (E,F)** 3,5 dpf: N=1,1 experiments; tmx2b^+/?^, n=9,6; tmx2b^-/-^, n=6,4. Two-way ANOVA, Tukey’s multiple comparisons test. **(G)** representative images of *mpeg1*:GFP+ microglia (green) and Lysotracker in the midbrain of *tmx2b*^*+/?*^ and *tmx2b*^*-/-*^ zebrafish at 3 and 5 dpf. Microglia in *tmx2b*^*-/-*^ zebrafish have an amoeboid morphology and higher lysosome concentration. Scale bar indicates 15 μm. **(H)** Quantification of lysosomal area within microglia in midbrain of *tmx2b*^*+/?*^ and *tmx2b*^*-/-*^ zebrafish at 3 and 5 dpf. Each dot represents the average value of six microglia from a single zebrafish brain. 3,5 dpf: N=1,1 experiment, *tmx2b*^*+/?*^ n=16,19, *tmx2b*^*-/-*^ n=9,13 zebrafish. Two-way ANOVA with Šídák›s multiple comparisons test. Data are represented as mean ± SD. *p < 0.05, **p < 0.01, ***p < 0.001, ****p < 0.0001.

Similarly, we assessed with specific markers oligodendrocyte precursor cells (OPCs) (*olig1:*NLS-mApple+) and myelination levels (*mbp:*EGFP-CAAX+). We observed decreased OPC numbers in *tmx2b*^*-/-*^ zebrafish compared to *tmx2b*^*+/?*^ at 3 and 5 dpf **(Figure 3C,D and Figure S10)**. Consistent with lower OPC numbers, myelination in the midbrain/hindbrain region was largely lacking in *tmx2b*^*-/-*^ larvae **(Figure 3E,F)**. The observed difference in OPCs appeared to be caused by a lack OPC proliferation between 2 and 3 dpf, as the total numbers of OPCs were similar at 2 and 3 dpf in *tmx2b*^*-/-*^ zebrafish **(Figure 3C,D)**.

Having shown effects on neurons, OPCs and myelination, we evaluated the effect of Tmx2b loss on microglia, brain-resident macrophages (Neutral Red, NR+ and *mpeg1:*EGFP+) (44). At 3 dpf microglia numbers were not different in *tmx2b*^*-/-*^ compared to *tmx2b*^*+/?*^, but they were more localized to the optic tecti regions and had an amoeboid, rounded morphology, indicative of a high level phagocytic activity **(Figure S11A-F)** (45). Increased lysotracker staining in microglia at 3 and 5 dpf is indeed consistent with highly phagocytic microglia **(Figure 3G,H)** (26). Microglia numbers were normal at 3 dpf but at 5 dpf microglia numbers had increased ∼2 fold compared to *tmx2b*^*+/-*^ **(Figure 3G,H and Figure S11D-F)**. Thus, *tmx2b*^*-/-*^ microglia are likely unaffected by Tmx2b loss as their initial development is normal. Furthermore, the microglia are capable of performing their physiological functions by proliferating as a response to neuronal cell death and actively phagocytizing neuronal cell debris (44).

### Neuronal cell death occurs rapidly at a specific time point in development

To obtain a better understanding of the onset of neuronal cell death in *tmx2b*^*-/-*^ zebrafish we performed overnight temperature controlled time-lapse imaging of the DsRed+ excitatory and GFP+ inhibitory neurons from 63 hpf till 75 hpf. Before onset of neuronal cell death the *tmx2b*^*-/-*^ brain developed normally and differentiated migrating neurons could be observed that were similar to those in *tmx2b+/ ?* zebrafish **(Video S10,11)**. Then neurons ceased to migrate within a 10 min time interval, and a brain deformation with loss of neurons was observed, with fluorescence intensity loss as a marker for neuronal cell loss over time. **(Figure 4A and Video S10,11)**. Onset of excitatory and inhibitory cells loss coincided and occurred within a ∼1.5-hour timeframe **(Figure 4B)**. Hence, cortical development appears entirely normal in *tmx2b*^*-/-*^ zebrafish until the onset neuronal cell death, which occurs suddenly and spans an ∼1.5 hours timeframe.

**Figure 4.**
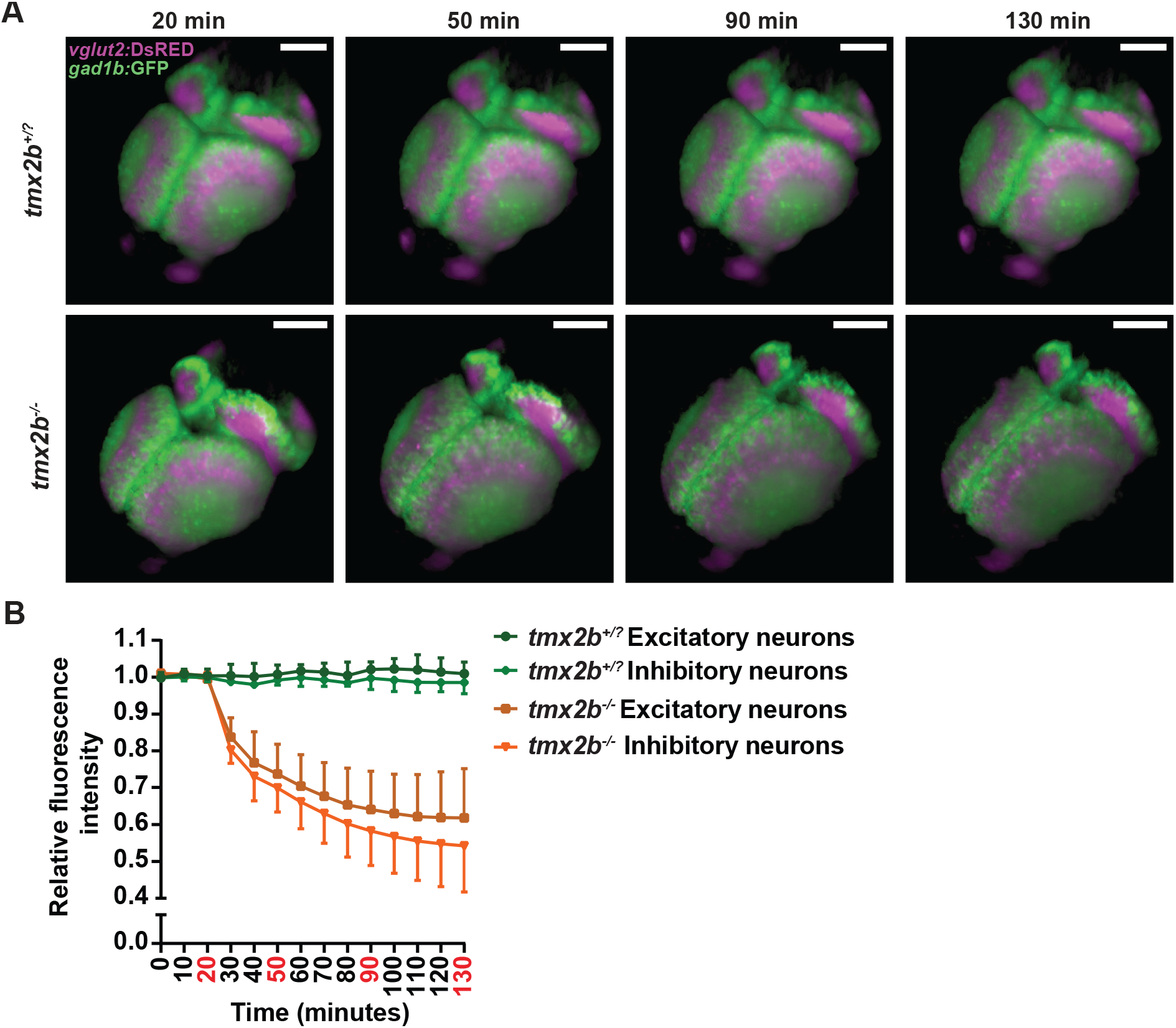
Neuronal cell death has a rapid onset. **(A)** 3D reconstructions of the *vglut2:*DsRED+ excitatory neurons (magenta) and *gad1b:*GFP+ inhibitory neurons (green) overnight imaging of *tmx2b*^*+/-*^ and *tmx2b*^*-/-*^. Zebrafish were imaged from 58 hpf till 75 hpf. This figure only shows the images around onset of the neuronal cell death in *tmx2b*^*-/-*^. **(B)** Quantification of relative fluorescence intensity of excitatory and inhibitory neurons in the midbrain as a measurement for neuronal cell loss. The neuronal cell loss starts within a 10 minute time period and the majority of cell loss occurs within ∼1.5 hour time window. n=3 zebrafish for both groups. Data are represented as mean ± SD.

### Modulating ROS does not affect the neuronal cell death

Since TMX2 has putative oxidoreductase activity and knockdown of TMX2 has been shown to increase ROS in cholangiocarcinoma cells *in vitro*, we reasoned that increased cellular ROS potentially could cause neuronal cell death (18, 46). Therefore, we tested the general antioxidant *N*-acetylcysteine (NAC) and two inhibitors of H_2_O_2_ synthesizing proteins in the ER (EN460: ERO1 inhibitor; GKT137831: NOX4 inhibitor) **(Table S3)** (47-50). All drugs did not ameliorate the *tmx2b*^*-/-*^ zebrafish phenotype, as all treated zebrafish still had growth delay and visible neuronal cell death **(Figure S12)**. To further investigate whether ROS contributed to the *tmx2b*^*-/-*^ phenotype, we increased cellular ROS by H_2_O_2_ treatment. Surprisingly, a decrease in visible brain necrosis was observed in *tmx2b*^*-/-*^ H_2_O_2_ treated zebrafish at 3 dpf **(Figure S13A-C)**. However, H_2_O_2_ treatment also resulted in a minor developmental delay in H_2_O_2_ treated *tmx2b*^+/-^, explaining the decrease in brain necrosis **(Figure S13B)**. Nonetheless, H_2_O_2_ treatment did not result in an earlier onset of the necrotic phenotype compared to untreated *tmx2b*^*-/-*^ zebrafish, suggesting that loss of neurons in *tmx2b*^-/-^ is not primarily driven by increased ROS.

### Ca^2+^ dysregulation in *tmx2b*^*-/-*^ zebrafish brain

Individuals with pathogenic *TMX2* variants often present with severe epilepsy (6, 7). It is unknown whether any epilepsy-induced neuronal cell damage and eventual death relates to the progressive disease course observed in these individuals (6, 7). Possibly, abnormal neuronal firing could contribute to neuronal cell death, and to test this hypothesis we treated zebrafish with the voltage-gated sodium channel blocker tricaine (51, 52). Strikingly, tricaine treatment prevented necrosis development in ∼66% of *tmx2b*^*-/-*^ zebrafish at 3 dpf **(Figure 5A-C)**. In contrast to untreated zebrafish, tricaine treated *tmx2b*^*-/-*^ zebrafish had a body length not different from treated and untreated control zebrafish; therefore, decreased necrosis could not be explained by a general developmental delay of the zebrafish **(Figure 4F)**. If inhibition of neuronal action potentials by anesthetic, tricaine, prevented necrosis, induction of neuronal activity or eliciting seizures could speed up the onset of necrosis . We utilized 4-Aminopyridine (AP), a potassium channel antagonist, to induce over-excitation of neurons in the *tmx2b*^*-/-*^ zebrafish, which did not show an earlier onset of neuronal cell death **(Figure S13D-F)** (53, 54). At 2 dpf all 4-AP treated *tmx2b*^*-/-*^ zebrafish showed normal development and displayed no gray discoloration in the brain, whereas at 3 dpf extensive necrosis, similar to the untreated group in the *txm2b*^*-/-*^ larvae, was observed **(Figure S13D-F)**. Altogether these data indicate that suppressing neuronal excitation, rescues neuronal cell death, whereas stimulating neuronal activity/seizures cannot accelerate the onset of neuronal cell death phenotype in *tmx2b*^*-/-*^ zebrafish. Alternatively, the effect of tricaine may be unrelated to seizure suppression and rather related to an additional function of voltage gated sodium channels in microglia activity (55).

**Figure 5.**
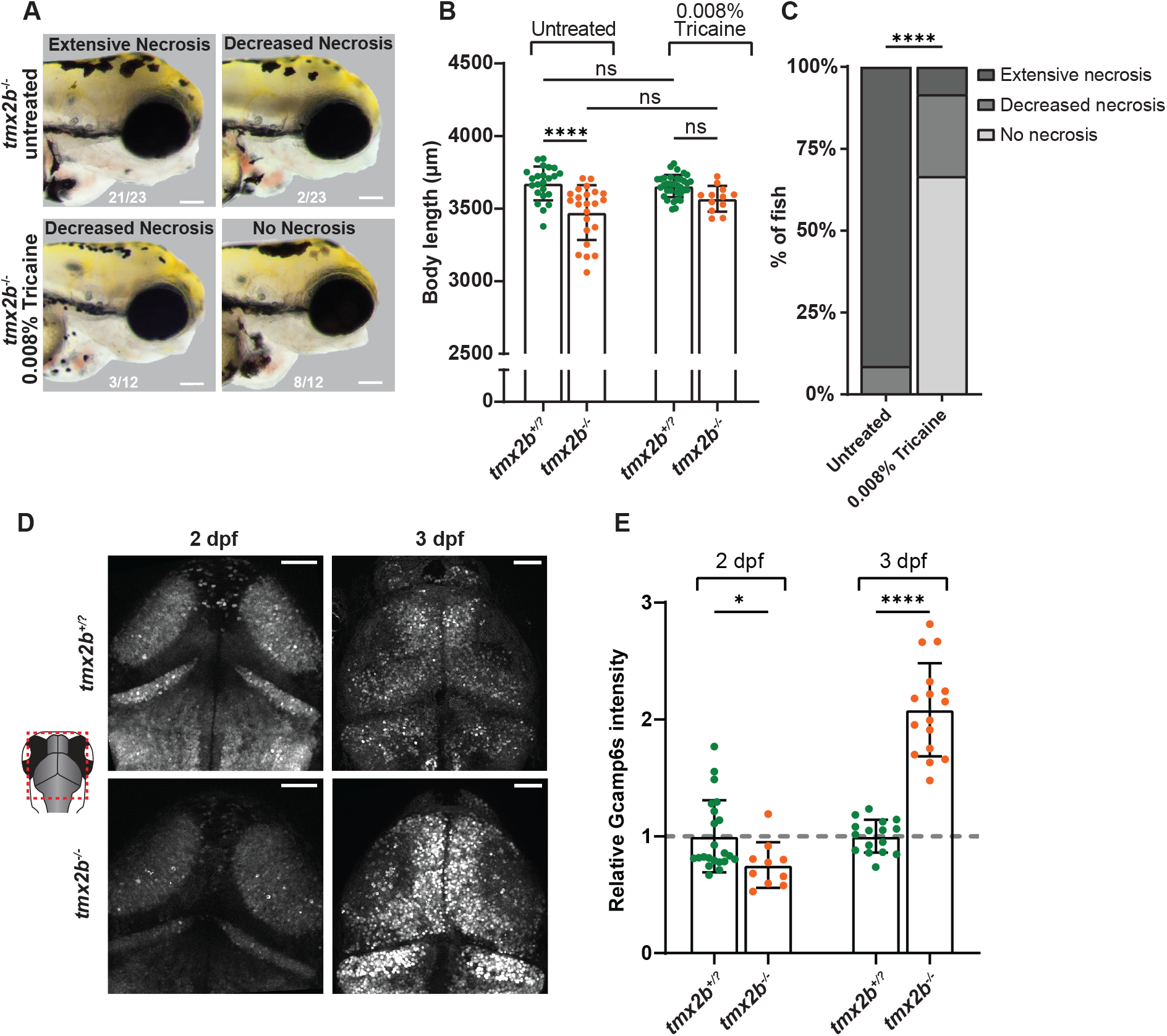
Ca^2+^ is dysregulated in neurons of *tmx2b*^*-/-*^ zebrafish. **(A)** Brightfield images lateral view head of 3 dpf zebrafish untreated (upper images) and 0.008% tricaine treated (lower images). Of the 0.008% tricaine treated *tmx2b*^*-/-*^ zebrafish >50% developed no necrosis at 3 dpf. Scale bar represents 100 μm. Bottom numbers indicate counts of zebrafish with specified genotype and phenotype. **(B)** Body length measurements of 3 dpf untreated and 0.008% tricaine treated *tmx2b*^*+/?*^ and *tmx2b*^*-/-*^ zebrafish. Two-way ANOVA with Tukey’s multiple comparisons test. Quantification of **(A)** Fisher’s exact test (extensive and decreased necrosis groups were combined for statistical test). **(B,C)** Untreated, 0.008% tricaine treated: N=2,2 experiments; *tmx2b*^*+/?*^, n=23,34 ; *tmx2b*^*-/-*^, n=23,12 zebrafish. **(D)** Representative images of *elavl3:*NLS-GCaMP6s neurons in *tmx2b*^*+/?*^ and *tmx2b*^*-/-*^ zebrafish brain at 2 and 3 dpf. Scale bars indicate 50 μm. **(B)** Relative NLS-GCaMP6s fluorescence intensity normalized to the average *tmx2b*^*+/?*^ value at 2 or 3 dpf. 2,3 dpf: N=1,1 experiment; *tmx2b*^*+/?*^, n=22,16; *tmx2b*^*-/-*^, n=10,16. Two-way ANOVA with Šídák›s multiple comparisons test on not normalized data. Data are represented as mean ± SD. *p < 0.05, **p < 0.01, ***p < 0.001, ****p < 0.0001.

Besides oxidoreductase activity, various PDIs also regulate Ca^2+^ flow between the ER and mitochondria (21). Since, TMX2 can localize to mitochondrial-ER contacts (MERCs) and can interact with calcium binding proteins (Vandervore 2019), we tested the hypothesis that Ca^2+^ homeostasis is dysregulated in *tmx2b*^*-/-*^ zebrafish (6, 19). We employed the neuronal Ca^2+^ sensor NLS-GCaMP6s (*elavl3:*NLS-GCaMP6s) in tricaine anesthetized zebrafish—thereby preventing any influence of neuronal activity—to assess baseline nuclear and cytosolic Ca^2+^ concentrations (34, 56). Already at 2 dpf, before onset of cell death, we observed that nuclear and cytoplasmic Ca^2+^ concentration in neurons were slightly decreased in *tmx2b*^-/-^ zebrafish compared to *tmx2b*^*+/?*^ **(Figure 5D, E and Figure S14)**. In contrast, at 3 dpf neuronal Ca^2+^ concentration was increased by ∼2-fold in *tmx2b*^*-/-*^ zebrafish **(Figure 5D, E and Figure S14)**. Our previous data could not identify abnormalities during the first 2 days of development in *tmx2b*^*-/-*^ zebrafish; however, these data suggest that Ca^2+^ dysregulation occurs prior onset of neuronal cell death.

## Discussion

Advanced genomic analysis has enabled identification of novel human brain disorders caused by genetic mutations in previously unknown genes at high pace. Hence, elucidation of the physiological gene function and disease mechanism follows identification of gene mutations and often requires recapitulation of the disorder in animal models. Here, we established a zebrafish disease model for TMX2-related malformation of cortical development and neurological disease and show that TMX2 is essential for survival of newborn neurons. We identified Tmx2b as the *TMX2* orthologue and found that *Tmx2b* inactivation interferes with zebrafish brain development. Both excitatory and inhibitory neurons in Tmx2b deficient zebrafish underwent cell death at the end of the zebrafish embryonic phase (39). This cell death had a sudden onset and occurred within a 1.5-hour timeframe and was not progressive till 5 dpf, beyond which age the embryos did not survive. Cell death occurred specifically in neurons, while OPCs were decreased in number without observable cell loss and myelination was largely absent in the brain. Neuronal progenitors/radial glial cells and microglia were not directly compromised by Tmx2b loss. Lastly, we showed that neuronal cell death could be suppressed by a voltage-gated sodium channel blocker and that Ca^2+^ dysregulation preceded neuronal loss, consistent with a role of TMX2 in regulating Ca^2+^ homeostasis in newborn neurons.

Individuals with biallelic pathogenic variants in *TMX2* present with severe NDD, epilepsy, primary and/or progressive microcephaly and polymicrogyria (6, 7). Primary microcephaly (i.e. microcephaly present at birth) results from either too little proliferation or increased cell death/apoptosis of the neural progenitor cells/radial glia cells (1, 57). Some individuals with *TMX2*-related brain malformation show progression of the microcephaly through infancy, which speaks for a progressive interference with brain development beyond an early proliferative stage. Our previous studies suggested that TMX2 loss results in cell death of the neural progenitor population (7). This study in zebrafish instead shows that not the neural progenitor cells, but the post-mitotic excitatory and inhibitory neurons undergo cell death at a specific point during brain development. The cell death of early born neurons observed in our zebrafish model could explain the microcephaly observed in individuals with pathogenic variants in *TMX2*, and how the neuronal damage could progress also after the initial proliferation stage.

The brain of newborns with *TMX2* loss often shows polymicrogyria, which is a defect of cortical organization, sometimes developmental, sometimes infectious, like for example the congenital cytomegalovirus (CMV) infection (6). Brain MRI and pathology of affected individuals showed that the polymicrogyria was unlayered, which is a sign of early disruption of cortical development between 13 and 16 gestational week, very similar to the CMV infection (6, 58, 59). In human brain affected by CMV infection, foci of necrosis close to the polymicrogyria region are detected, suggesting that localized cell death is linked to the polymicrogyria (59, 60). We hypothesize that the necrotic and apoptotic neurons during brain development in the *tmx2b*^*-/-*^ zebrafish recapitulate events leading to the cortical malformation seen on MRI and pathology of individuals with pathogenic *TMX2* variants. Our findings and conclusions also support that a damaging process resulting in neuronal cell death in the cortical plate is causal for the MCD.

The phenotype of *tmx2b-/* zebrafish appears more severe than what is typically observed in individuals with pathogenic *TMX2* variants (6, 7). While some affected individuals with pathogenic *TMX2* variants do not survive beyond the first month of life, others can reach adulthood, unlike the *tmx2b*^*-/-*^ zebrafish (6). Although *TMX2* pathogenic variants are supposed to have a loss-of-function effect, complete absence of TMX2 has not been proven in all affected individuals with biallelic variants (6, 7). Most affected individuals present either with biallelic missense variants or with compound heterozygous truncating, i.e. null mutation, and a missense variant. *Tmx2* knockout mice are embryonically lethal and our *tmx2b*^*-/-*^ zebrafish also do not survive beyond 5 dpf (61). This suggests that a complete loss of *TMX2* is lethal during embryonic development and could explain why no individuals have been described until now with biallelic truncating variants.

Although *in vitro* studies indicated that loss of TMX2 increases cellular ROS, we were unable to modulate the phenotype of *tmx2b*^*-/-*^ zebrafish by either increasing or decreasing ROS (18). Another possible explanation for the neuronal cell death is that Tmx2b loss results in a severe status epilepticus, which is known to be detrimental for neurons (62). By anesthetizing our *tmx2b*^*-/-*^ we could prevent neuronal cell death onset. However, drug-induced epilepsy did not result in an earlier onset of the phenotype, suggesting that prolonged and excessive neuronal activity is not the only cause of the *tmx2b*^*-/-*^ cell death, but could still be its consequence. Our results, however, point towards a general Ca^2+^ dysregulation in neurons. Ca^2+^ concentrations in the nucleus, which are indirectly a measurement for Ca^2+^ in the cytosol, were decreased at 2 dpf and increased at 3 dpf in *tmx2b*^*-/-*^ zebrafish (56). Tricaine anesthesia diminishes neuronal action potentials, consequently inhibiting Ca^2+^ influx in neurons, which could counteract the Ca^2+^ dyshomeostasis in *tmx2b*^*-/-*^ zebrafish brain.

Intracellular Ca^2+^ functions as a second messenger in various cellular processes including neurotransmitter release and the ER serves as the largest Ca^2+^ storing organelle (63, 64). The ER contains various Ca^2+^ channels that transport Ca^2+^ into the cytosol and subsequently towards the mitochondria. Ca^2+^ transport from the ER towards the mitochondria increases oxidative phosphorylation, but prolonged Ca^2+^ flux can also induce apoptosis (21). Ca^2+^ is released from the ER via the ryanodine receptors (RyR) and inositol 1,4,5-trisphosphate receptors (IP3Rs) of which the IP3R1 and RyR3 are primarily utilized in the brain (65, 66). Loss-of-function variants in *ITPR3*, encoding IP3R1, are associated with spinocerebellar ataxia 15 and knockout mice also display a cerebellar ataxia phenotype (67). Since we observe a brain specific effect of Tmx2b loss, without cerebellar anomalies, it seems unlikely that loss of TMX2 directly impairs the function of IP3R1. Ca^2+^ uptake from the cytosol towards the ER is executed by ATP2A2/SERCA2, which is known interactor of TMX2 (6). Heterozygous loss-of-function variants in *ATP2A2* cause Darier disease, which is primarily a skin disorder; however, neuropsychiatric diseases, such as bipolar disorder and schizophrenia, are also associated with this disease (68, 69). Interestingly, brain specific *Atp2a2* knockout in mice is embryonically lethal and display destructive intracerebral hemorrhages (70). Whether these intracerebral hemorrhages were the result of a primary vascular defect or secondary to neuronal cell death was not determined (70). Lack of physiological interaction with Ca^2+^ regulating proteins could therefore be the cause of neuronal cell death in the absence of Tmx2b (71, 72).

A previous study showed that treatment of zebrafish with either rotenone and azide— a mitochondrial complex I and IV inhibitor, respectively— also induced cell death exclusively in the brain, reminiscent of our observations in the *tmx2b*^-/-^ zebrafish (41). Fibroblasts of individuals with pathogenic variants in *TMX2* also display mitochondrial dysfunction and specifically suppressed mitochondrial respiration after treatment with mitochondrial uncoupler FCCP, which reflects reduced reserve capacity (6). Additionally, all tested cell strains showed decreased rotenone-dependent respiration, which indicates reduced activity of complex I (6). Altogether, this points towards a potential mitochondrial impairment in the *tmx2b*^*-/-*^ zebrafish. As mitochondrial respiration is regulated by Ca^2+^ flux from the ER to mitochondria via MERCs and TMX2 is localized at these contact points, loss of TMX2 could compromise this Ca^2+^ flow, consequently impairing mitochondrial function. This mitochondrial dysfunction results in neuronal cell death specifically, due to the relatively high energy demand of newborn neurons, required for migration, axon and dendritic growth, and synapse formation.

Although TMX2 is ubiquitously expressed we only observed a largely neuron specific effect in *tmx2b*^*-/-*^ zebrafish, and a reduction of OPCs and myelination (8). Between 2 and 3 dpf OPC numbers remained the same in *tmx2b*^*-/-*^ zebrafish, suggesting a lack of proliferation and not necessarily cell death of OPCs. It is well established that OPC proliferation and myelination is dependent on neuronal activity and the presence of axons (30, 35, 73, 74). Therefore, the decreased OPC numbers and lack of myelination is either a secondary effect to the neuronal cell death, or the result of insufficient differentiation of common radial glial progenitors into OPC. We did not detect similar extent of cell death in any other organ than the brain at 3 dpf to 5 dpf. Hence, cell death caused by Tmx2b loss appears limited to neurons. Mitochondrial dysfunction in *tmx2b*^*-/-*^ in combination with a higher energy demand of neurons provides a possible explanation for the neuronal cell death. Ca^2+^ signaling and currents differ between neurons and glia cells, and also between various neuronal types. Therefore, an alternative explanation for the neuronal cell death in the central brain regions of *tmx2b*^*-/-*^ zebrafish could be dependent on how Ca^2+^ signaling is organized in the different cell types throughout the central nervous system (75).

This study has certain limitations to acknowledge. Firstly, we only conducted experiments on germ line full body *tmx2b* knockouts. An inducible knockout approach could determine whether *tmx2b* is only required for neuronal survival between 2 and 3 dpf (76). If Tmx2b is only essential for neurons during this time period, knocking out Tmx2b after 3 dpf would not cause neuronal cell death and *tmx2b*^*-/-*^ zebrafish may survive into adulthood. Furthermore, conditional *tmx2b* knockout of the different brain cell types could be beneficial to determine if malfunction of glia cell types is in part responsible for the neuronal cell death. The second limitation of our study is that we did not confirm the effect of our ROS reducing drugs with a cellular ROS specific staining. We tried to visualize cellular ROS with 2′,7′-dichlorofluorescein diacetate (DCFH-DA; Sigma, D6883), which has been used previously *in vivo* in zebrafish (77, 78). We noticed that the tissue penetrance of DCFH-DA *in vivo* was very limited, and staining was mainly observed in either the larval gut or blood vessels, similar to previous studies (77, 78). One study was able to detect ROS punctae with DCFH-DA in the spinal cord region at 3.5 dpf, but we were unable to recapitulate this observation (77).

In conclusion, we established, using a zebrafish genetic disease model, that Tmx2b is essential for survival of specifically newborn neurons. Loss of Tmx2b does not affect initial brain and neuronal development; however, at the end of the embryonic developmental phase excitatory and inhibitory neurons undergo cell death within a 1.5-hour timeframe. The cause of the contemporary decrease of normal oligodendrocyte population and insufficient myelination needs to be further explored. Radial glial cell and microglial populations are not affected by Tmx2b loss. Based on these observations we hypothesize that TMX2 is essential for the survival of post-mitotic newborn neurons during brain development. Its loss causes neuronal dysregulation of Ca^2+^, which through impaired mitochondrial function, could cause cell death and depletion of the newborn neuron population, providing a model for the microcephaly and disruptive polymicrogyria observed in individuals with *TMX2* mutations.

## Supporting information

Video S4 tmx2b ko

Video S5 tmx2b het

Video S6 3 dpf tmx2b ko

Video S7 3 dpf tmx2b ko delayed response

Video S8 4 dpf tmx2b het

Video S9 4 dpf tmx2b ko

Video S10 3D representation tmx2b het

Video S11 3D representation tmx2b ko

Video S1 1 dpf tmx2b het

Video S2 1 dpf tmx2b ko

Video S3 2 dpf tmx2b het

## Acknowledgements

The authors would like to thank prof. dr. Thomas Simmen for the fruitful discussions on TMX2.

