## Supplementary material for "The non-canonical thioreductase TMX2 is essential for neuronal survival during embryonic brain development": Suppl Figures 1-14

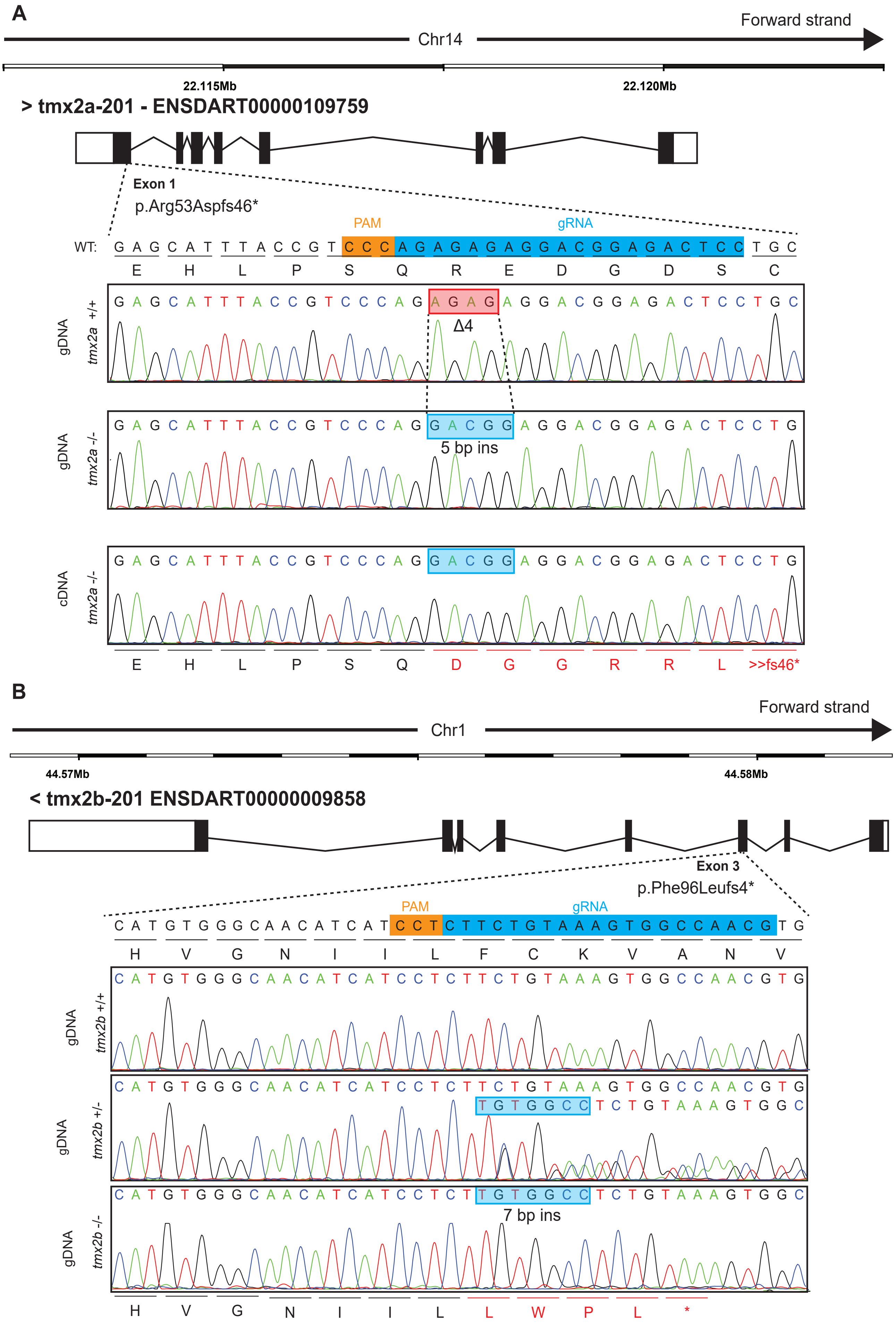


**Figure S1. *tmx2a* and *tmx2b* mutagenesis with CRISPR-Cas9. (A)** Schematic representation of the *tmx2a* gene (Ensembl transcript ID:ENSDART00000109759.5) the gRNA was designed to target exon 1. The adult *tmx2a*$^{-/-}$ zebrafish had a homozygous c.157_160delinsGACGG, p.Arg53Aspfs46* mutation. **(B)** Schematic representation of the *tmx2b* gene (Ensembl transcript ID:ENSDART00000009858.6). the gRNA was designed to target exon 3. The adult *tmx2b*$^{-/-}$ zebrafish had a heterozygous c.285_286insTGTGGCC, p.Phe96Leufs4* mutation.

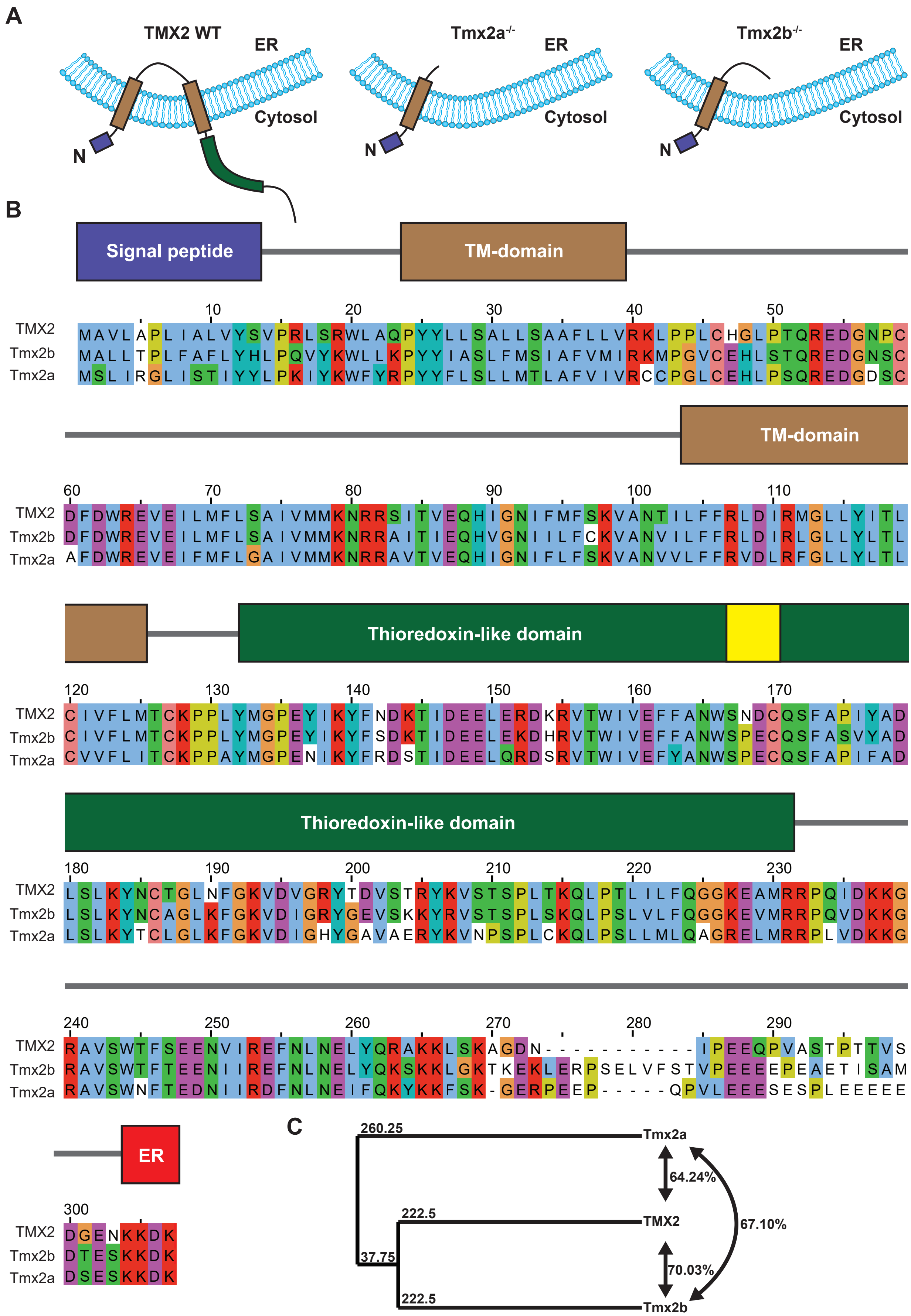


**Figure S2. TMX2, Tmx2a and Tmx2b protein comparison.** **(A)** Schematic representation of TMX2 orientation in the ER membrane. The potential proteins generated from the mutant alleles only contain the N-terminal signal peptide and first transmembrane domain and lack the catalytic domain of the protein. **(B)** Protein alignment of Tmx2, Tmx2a and Tmx2b by the Jalview v2.11.2.7 software. Color coding is according to the Clustal algorithm. The protein domains drawn above the protein sequence are based on the human TMX2 protein (Interpro ID: Q9Y320). The yellow box indicates the catalytic S-X-X-C site of TMX2 **(C)** Average distance tree of the BLOSUM62 algorithm, indicating that the Tmx2b is closer related to TMX2 than Tmx2a. Percentages right side indicate protein sequence similarity. Abbreviations: ER, ER retention signal; TM, transmembrane domain.

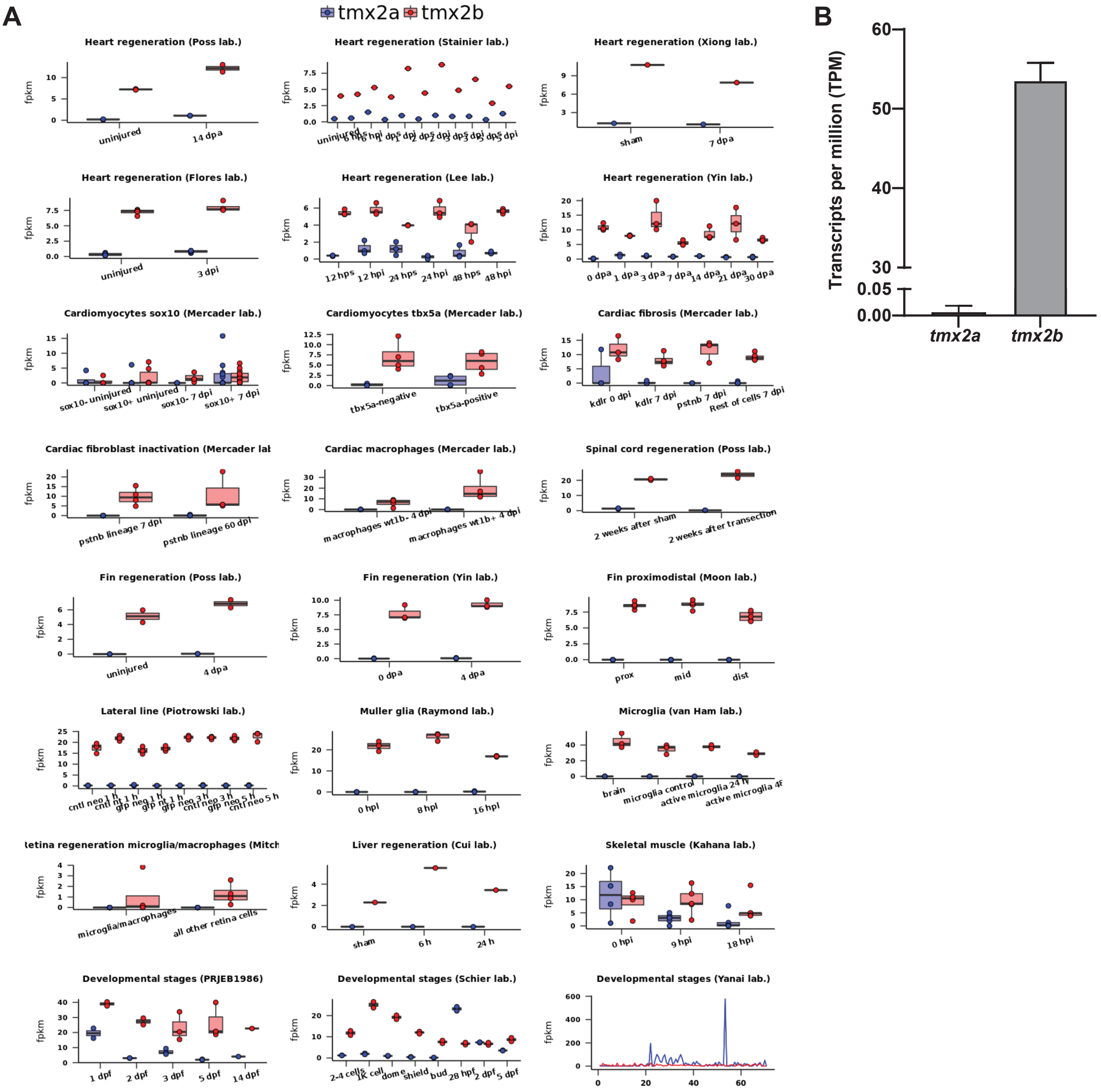


**Figure S3. Tmx2b is the homologue of TMX2. (A)** Expression level plots of *tmx2a* and *tmx2b* from different RNA-seq data-sets (source: http://zfregeneration.org/). all different data-sets show that *tmx2b* is the mainly expressed gene. **(B)** In house RNA-seq data of WT zebrafish brain at 5 days post fertilization (dpf) showing that *tmx2a* has no expression and *tmx2b* has an average expression > 50 TPM (n=3, each sample is a pool of brain (sample 1 = 16 brains, sample 2 = 22 brains and sample 3 = 25 brains)

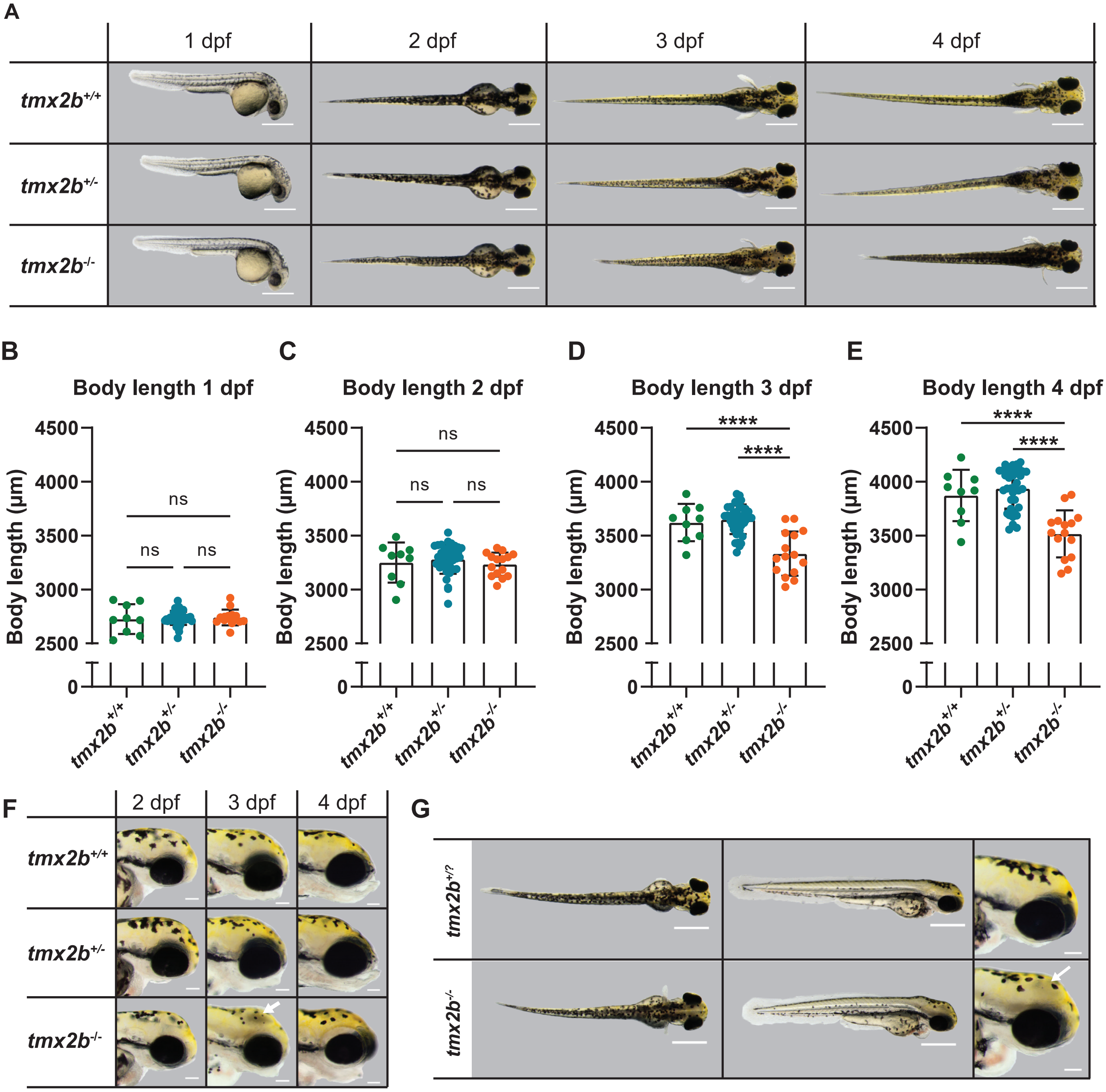


**Figure S4.** ***tmx2b*$^{-/-}$ zebrafish display a developmental decline from 3 dpf onwards.** **(A)** Representative images of same *tmx2b*$^{+/+}$, *tmx2b*$^{+/-}$ and *tmx2b*$^{-/-}$ zebrafish from 1 to 4 days post fertilization (dpf). *tmx2b*$^{-/-}$ have normal body morphology till 2 dpf and from 3 dpf onwards display a developmental decline. Scale bars indicate 500 µm. **(B,C,D,E)** Body length measurements of *tmx2b*$^{+/+}$, *tmx2b*$^{+/-}$ and *tmx2b*$^{-/-}$ zebrafish from 1 till 4 dpf. t*mx2b*$^{-/-}$ zebrafish have a smaller body length indicative for a developmental delay from 3 dpf onwards. N=2 experiments, t*mx2b*$^{+/+}$ n=9, t*mx2b*$^{+/-}$ n=37, t*mx2b*$^{-/-}$ n=15 zebrafish, one-way ANOVA with Tukey's multiple comparisons test. **(F)** Representative images of *tmx2b*$^{+/+}$, *tmx2b*$^{+/-}$ and *tmx2b*$^{-/-}$ zebrafish brightfield images lateral view of head. *tmx2b*$^{-/-}$ zebrafish develop a gray discoloration in brain region (white arrow) indicative of necrosis. This gray discoloration is not always present at 4 dpf in tmx2b-/- zebrafish. Scale bars indicate 100 µm **(G)** Images showing the onset of visible necrosis (white arrow) in the brain of t*mx2b*$^{-/-}$ zebrafish. Onset of the brain necrosis occurs between the pec-fin stage (60 hpf) and protruding-mouth stage (72 hpf); e.g. embryonic to larval transition.Scale bars whole fish indicate 500 µm. Scale bars fish head indicate 100 µm. Data are represented as mean ± SD. *$p < 0.05$, **$p < 0.01$, ***$p < 0.001$, ****$p < 0.0001$.

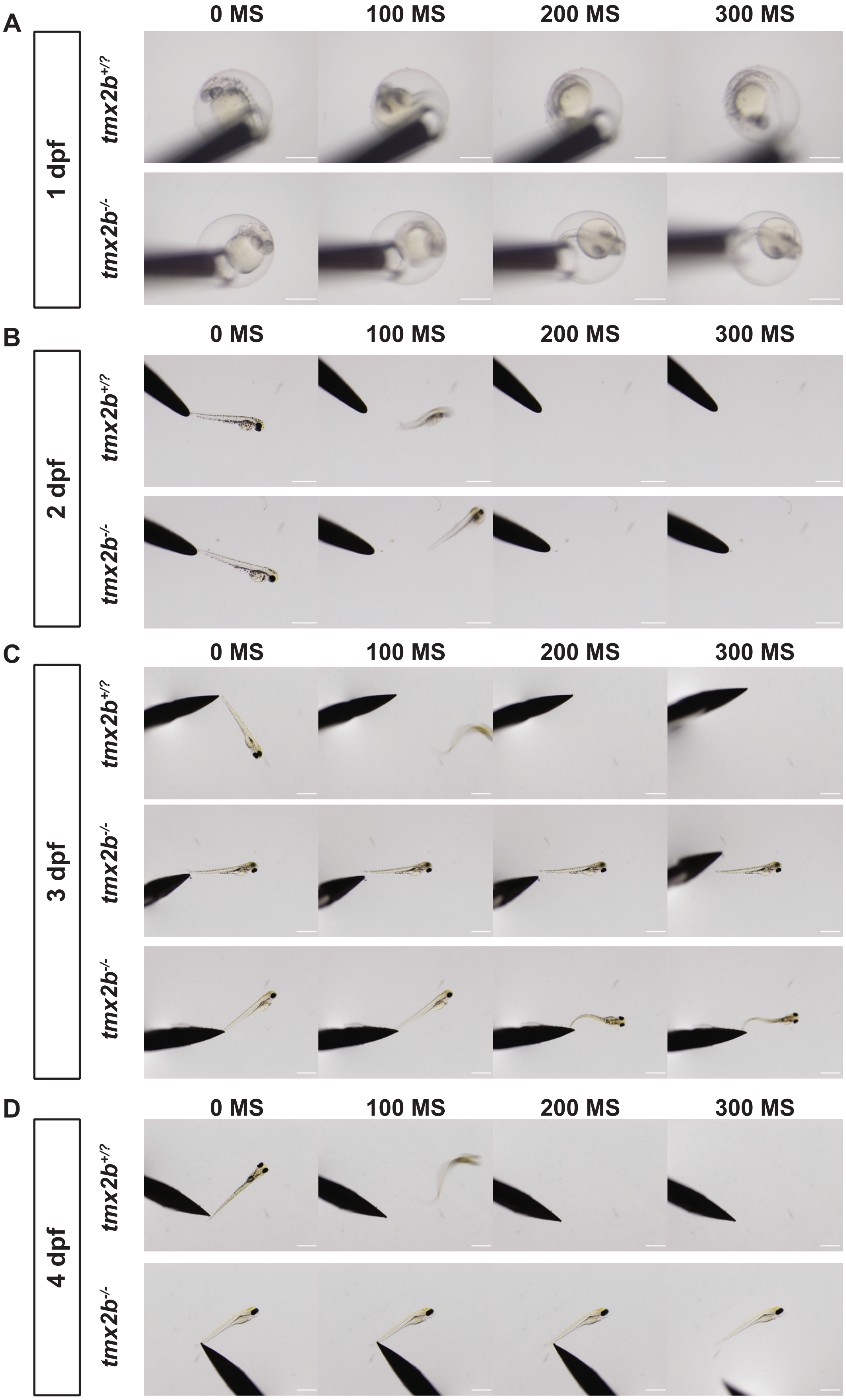


**Figure S5. Touch response assay from 1 till 4 dpf. (A)** Touch response at 1 dpf. Touch response was normal if the zebrafish had an immediate twitching movement upon touch. Both t*mx2b*$^{+/?}$ and t*mx2b*$^{-/-}$ zebrafish display a normal touch response at 1 dpf. **(B)** Touch response at 2 dpf. Both t*mx2b*$^{+/?}$ and t*mx2b*$^{-/-}$ zebrafish display a normal touch response and immediately swim away upon touch by needle. **(C)** Touch response at 3 dpf. t*mx2b*$^{+/?}$ zebrafish immediately swim away upon touch by needle. t*mx2b*$^{-/-}$ zebrafish no longer swim away upon touch. Middle panel shows a zebrafish with no movements and lower panel a fish with ineffective movements (classified under delayed respose). **(D)** Touch response at 4 dpf. Similar to 3 dpf, t*mx2b*$^{+/?}$ zebfrafish swims away upon touch and t*mx2b*$^{-/-}$ zebrafish are unable to move. N=2 experiments, t*mx2b*$^{+/?}$ n=58, t*mx2b*$^{-/-}$ n=27 zebrafish. Scale bars 1 dpf indicate 500 µm. Scale bars 2-4 dpf indicates 1000 µm.

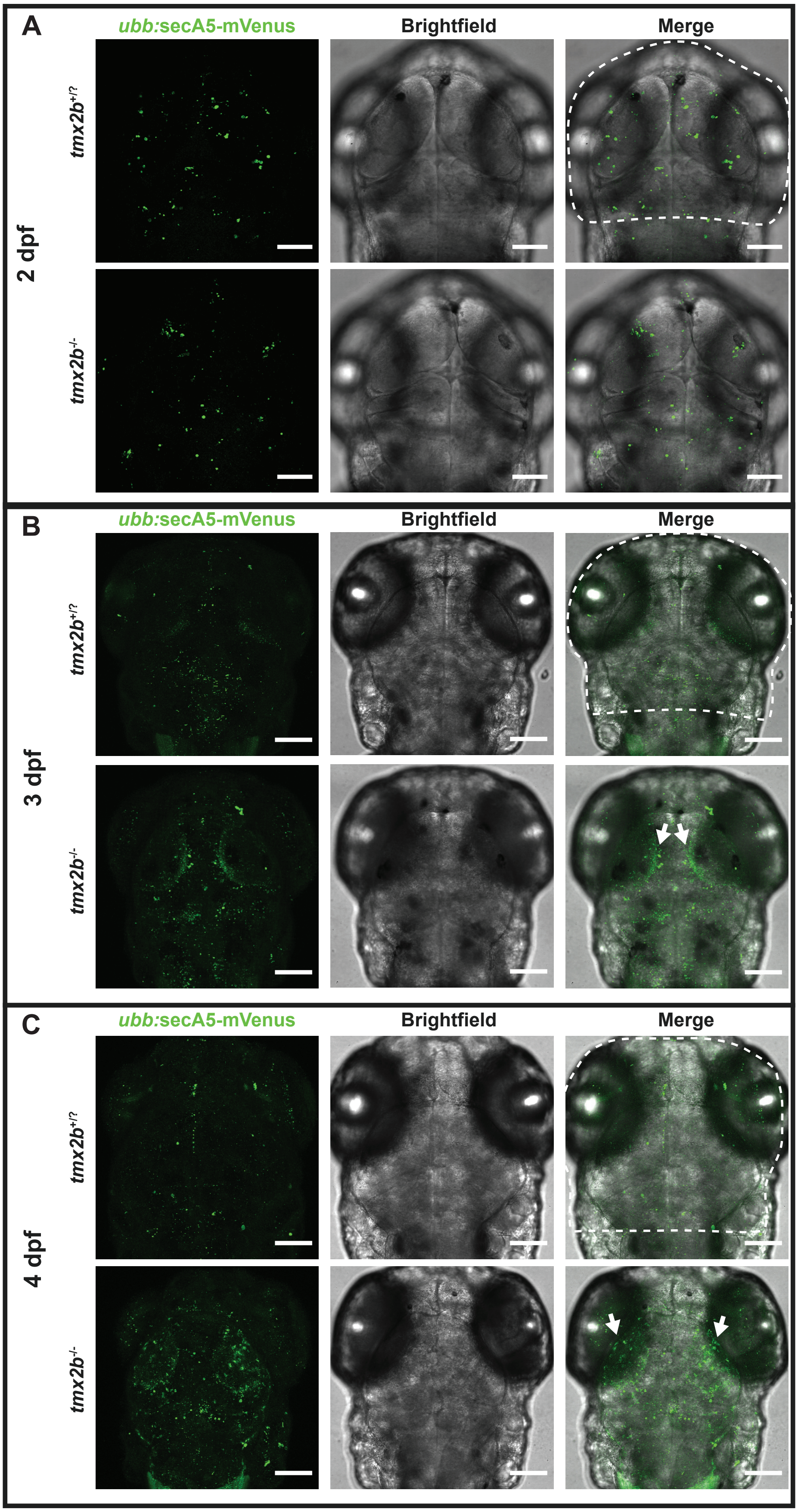


**Figure S6. Apoptotic clusters are increased in *tmx2b*$^{-/-}$ brain at 3 and 4 dpf (A,B,C)** Representative images of *ubb:*secA5-mVenus+ (green) apoptotic clusters in *tmx2b*$^{+/?}$ and t*mx2b*$^{-/-}$ zebrafish at 2, 3 and 4 dpf. Dashed lines in merged images indicate brain area where the apoptotic clusters were counted. t*mx2b*$^{-/-}$ zebrafish have an increased number of apoptotic clusters in the brain more pronounced in the optic tecti (white arrows) at 3 and 4 dpf. 2 dpf: N=2 experiments, t*mx2b*$^{+/?}$ n=30, t*mx2b*$^{-/-}$ n=6 zebrafish. 3 dpf: N=2 experiments, t*mx2b*$^{+/?}$ n=30, t*mx2b*$^{-/-}$ n=9 zebrafish. 4 dpf: N=1 experiment, t*mx2b*$^{+/?}$ n=11, t*mx2b*$^{-/-}$ n=6 zebrafish. Scale bars 2 dpf indicate 75 μm. Scale bars 3 dpf indicate 100 μm.

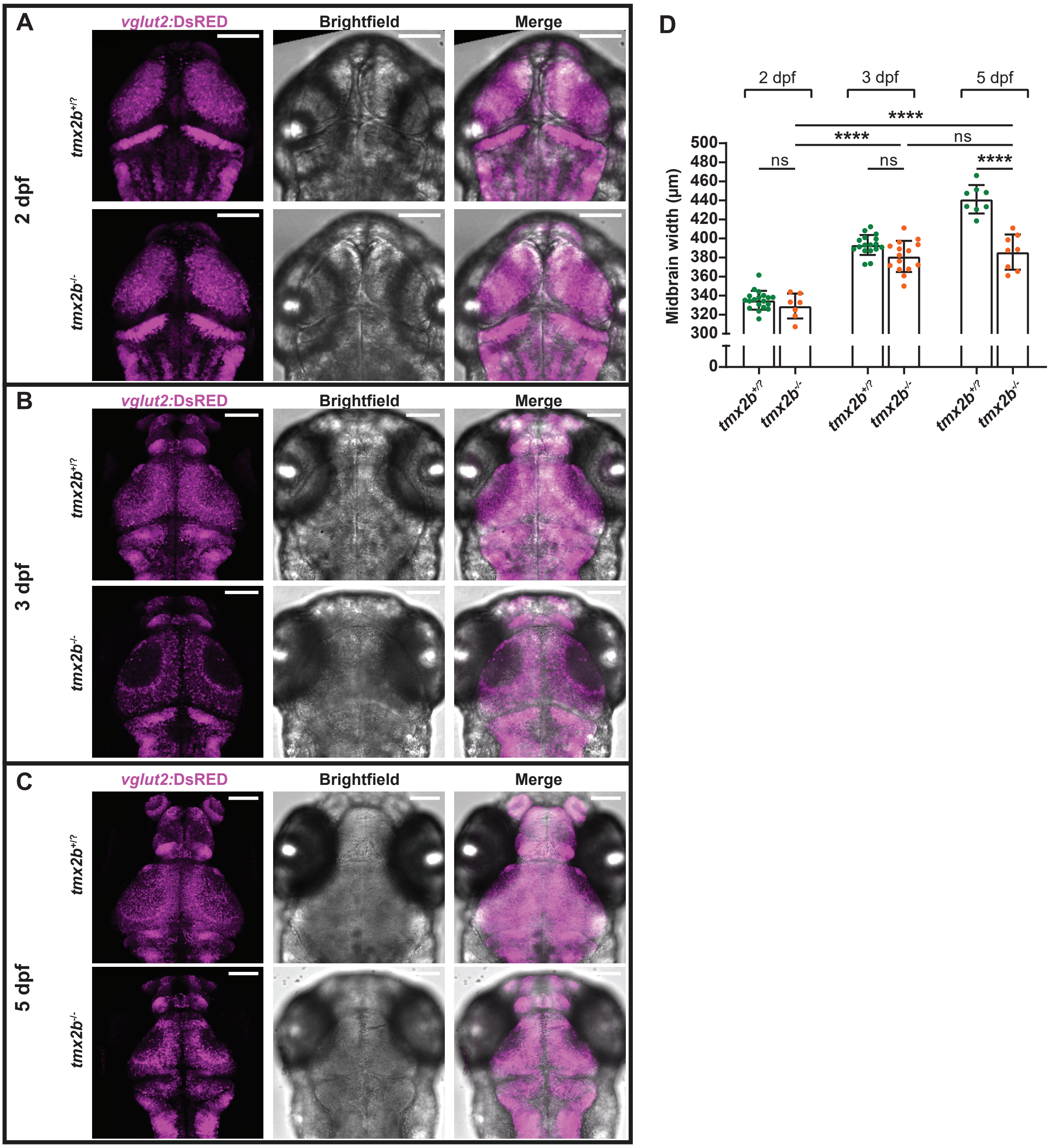


**Figure S7. Excitatory neurons in *tmx2b*$^{-/-}$ zebrafish brain undergo cell death between 2 and 3 dpf. (A,B,C)** Representative images of *vglut2*:DsRED+ (magenta) excitatory neurons in t*mx2b*$^{+/?}$ and t*mx2b*$^{-/-}$ zebrafish at 2, 3 and 5 dpf. t*mx2b*$^{-/-}$ zebrafish display excitatory neuronal cell loss at 3 dpf. Scale bars indicate 100 µm. **(D)** Midbrain width measurements t*mx2b*$^{+/?}$ and t*mx2b*$^{-/-}$ zebrafish at 2, 3 and 5 dpf. The largest diameter of the excitatory neurons was measured for the midbrain width. 2 dpf: N=1 experiment, t*mx2b*$^{+/?}$ n=19, t*mx2b*$^{-/-}$ n=7 zebrafish. 3 dpf: N=2 experiments, t*mx2b*$^{+/?}$ n=17, t*mx2b*$^{-/-}$ n=15 zebrafish. 5 dpf: N=1 experiment, t*mx2b*$^{+/?}$ n=8, t*mx2b*$^{-/-}$ n=8 zebrafish. Two-way ANOVA, Tukey's multiple comparisons test. Data are represented as mean ± SD. *$p < 0.05$, **$p < 0.01$, ***$p < 0.001$, ****$p <$ 0.0001.

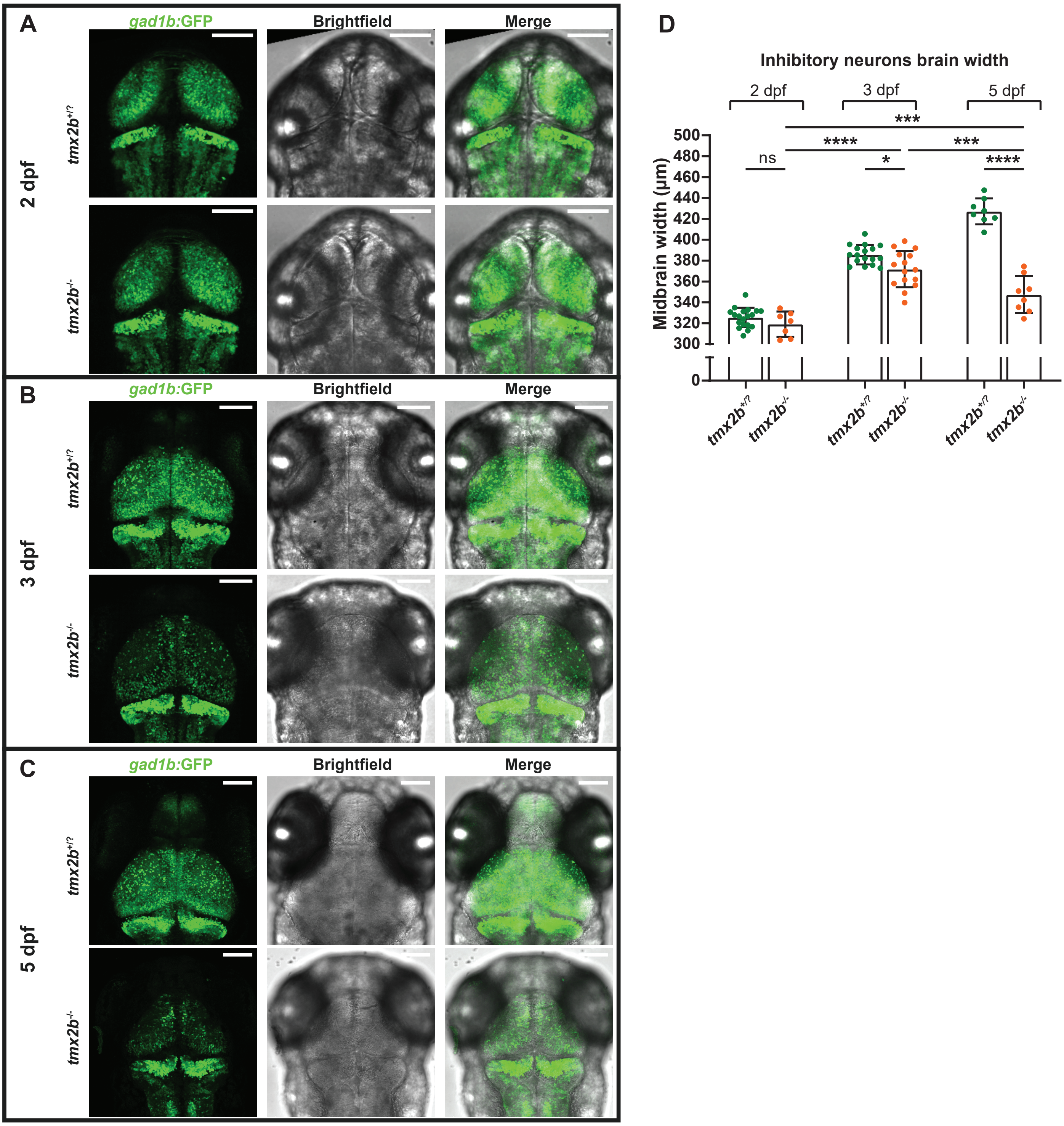


**Figure S8. Inhibitory neurons in in *tmx2b*$^{-/-}$ zebrafish brain undergo cell death between 2 and 3 dpf. (A,B,C)** Representative images of *gad1b:*GFP+ (green) inhibitory neurons in t*mx2b*$^{+/?}$ and *tmx2b*$^{-/-}$ zebrafish at 2, 3 and 5 dpf. t*mx2b*$^{-/-}$ zebrafish display inhibitory neuronal cell loss at 3 dpf. Scale bars indicate 100 µm. **(D)** Midbrain width measurements t*mx2b*$^{+/?}$ and t*mx2b*$^{-/-}$ zebrafish at 2, 3 and 5 dpf. The largest diameter of the inhibitory neurons was measured for the midbrain width. 2 dpf: N=1 experiment, t*mx2b*$^{+/?}$ n=19, t*mx2b*$^{-/-}$ n=7 zebrafish. 3 dpf: N=2 experiments, t*mx2b*$^{+/?}$ n=17, t*mx2b*$^{-/-}$ n=15 zebrafish. 5 dpf: N=1 experiment, t*mx2b*$^{+/?}$ n=8, t*mx2b*$^{-/-}$ n=8 zebrafish. Two-way ANOVA, Tukey's multiple comparisons test. *$p < 0.05$, ***$p < 0.001$, ****$p < 0.0001$.

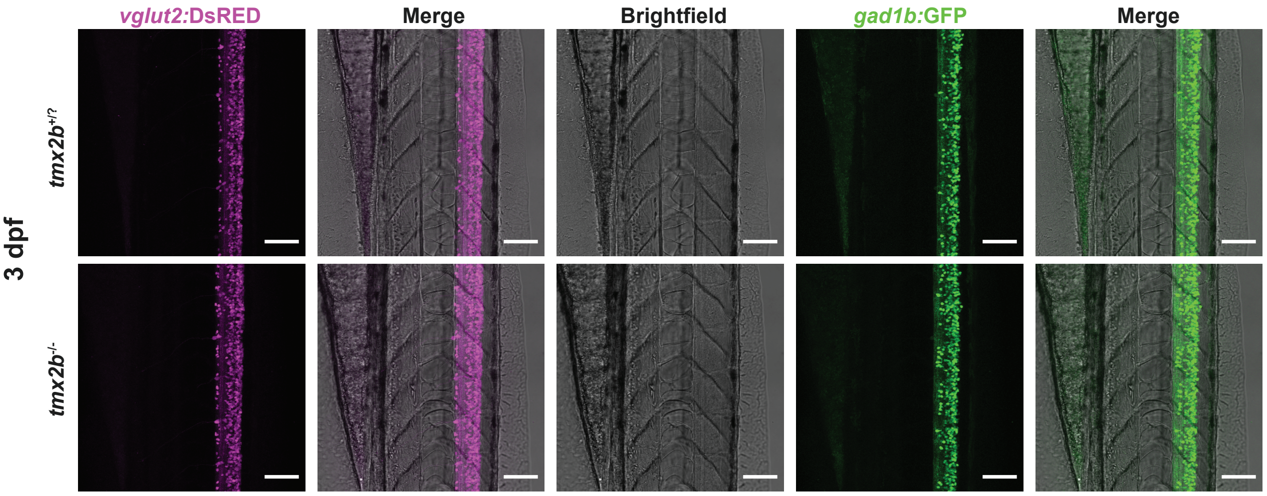


**Figure S9. Excitatory and inhibitory neurons in in *tmx2b*$^{-/-}$ zebrafish spinal cord are unaffected.** Representative images of *vglut2:*DsRED+ (magenta) excitatory and *gad1b:*GFP+ (green) inhibitory neurons in spinal cord in t*mx2b*$^{+/?}$ and t*mx2b*$^{-/-}$ zebrafish at 3 dpf. Contrary to the central brain regions the neurons in the spinal cord are unaffected by Tmx2 loss. N=2 experiments, t*mx2b*$^{+/?}$ n=15, t*mx2b*$^{-/-}$ n=10 zebrafish. Scale bars indicate 75 µm.

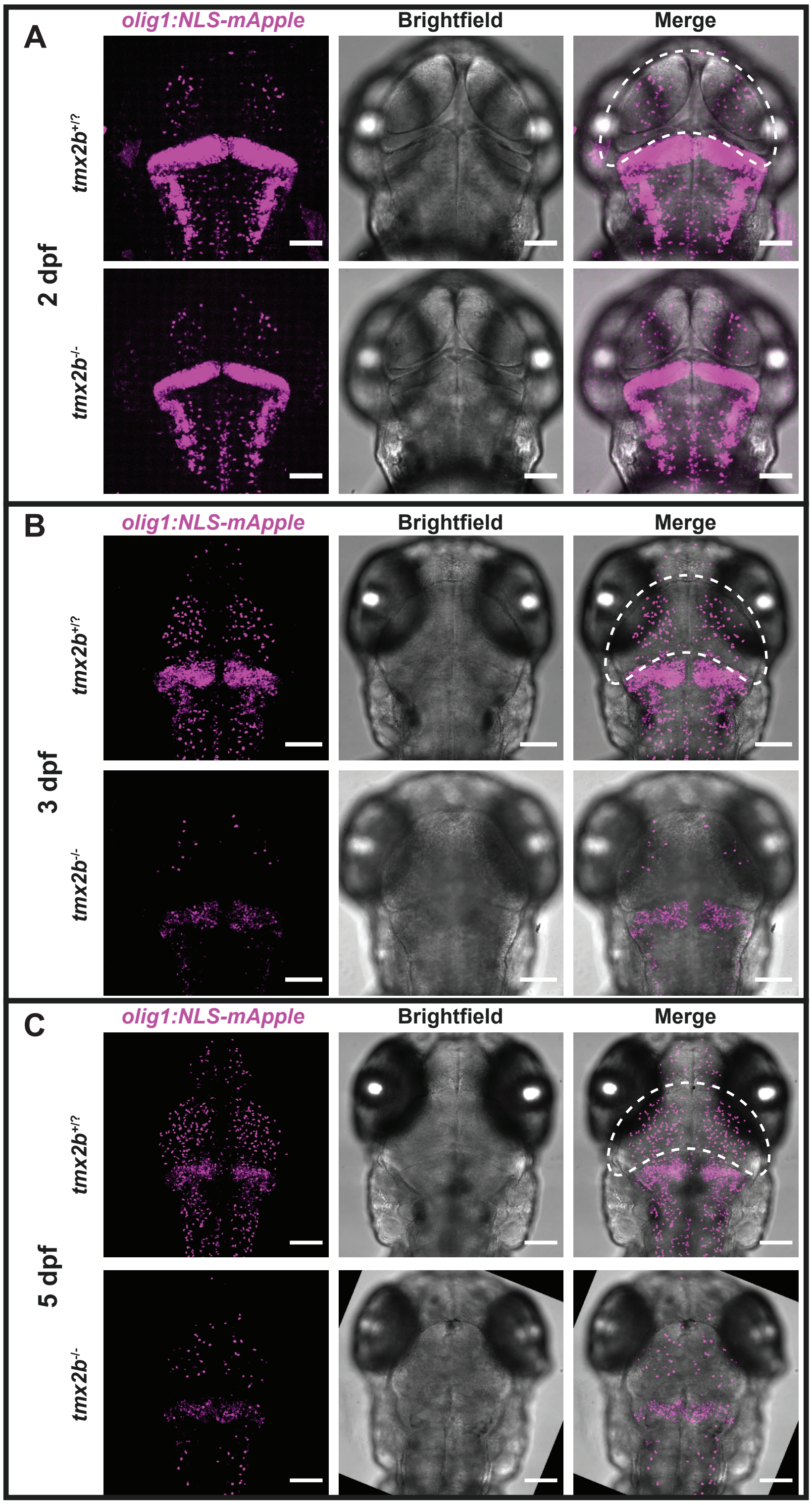


**Figure S10. Oligodendrocyte precursor cells (OPC) are decreased in *tmx2b*$^{-/-}$ zebrafish brain. (A,B,C)** Representative images of *olig1:*NLS-mApple (magenta) OPCs in t*mx2b*$^{+/?}$ and t*mx2b*$^{-/-}$ zebrafish at 2, 3 and 5 dpf. Scale bars 2 dpf indicate 75 μm. Dashed line indicates measured area. Scale bars 3 dpf indicates 100 μ m. 2 dpf: N=1 experiment, t*mx2b*$^{+/?}$ n=13, t*mx2b*$^{-/-}$ n=3 zebrafish. 3 dpf: N=1 experiment, t*mx2b*$^{+/?}$ n=9, t*mx2b*$^{-/-}$ n=6 zebrafish. 5 dpf: N=1 experiment, t*mx2b*$^{+/?}$ n=7, t*mx2b*$^{-/-}$ n=3 zebrafish.

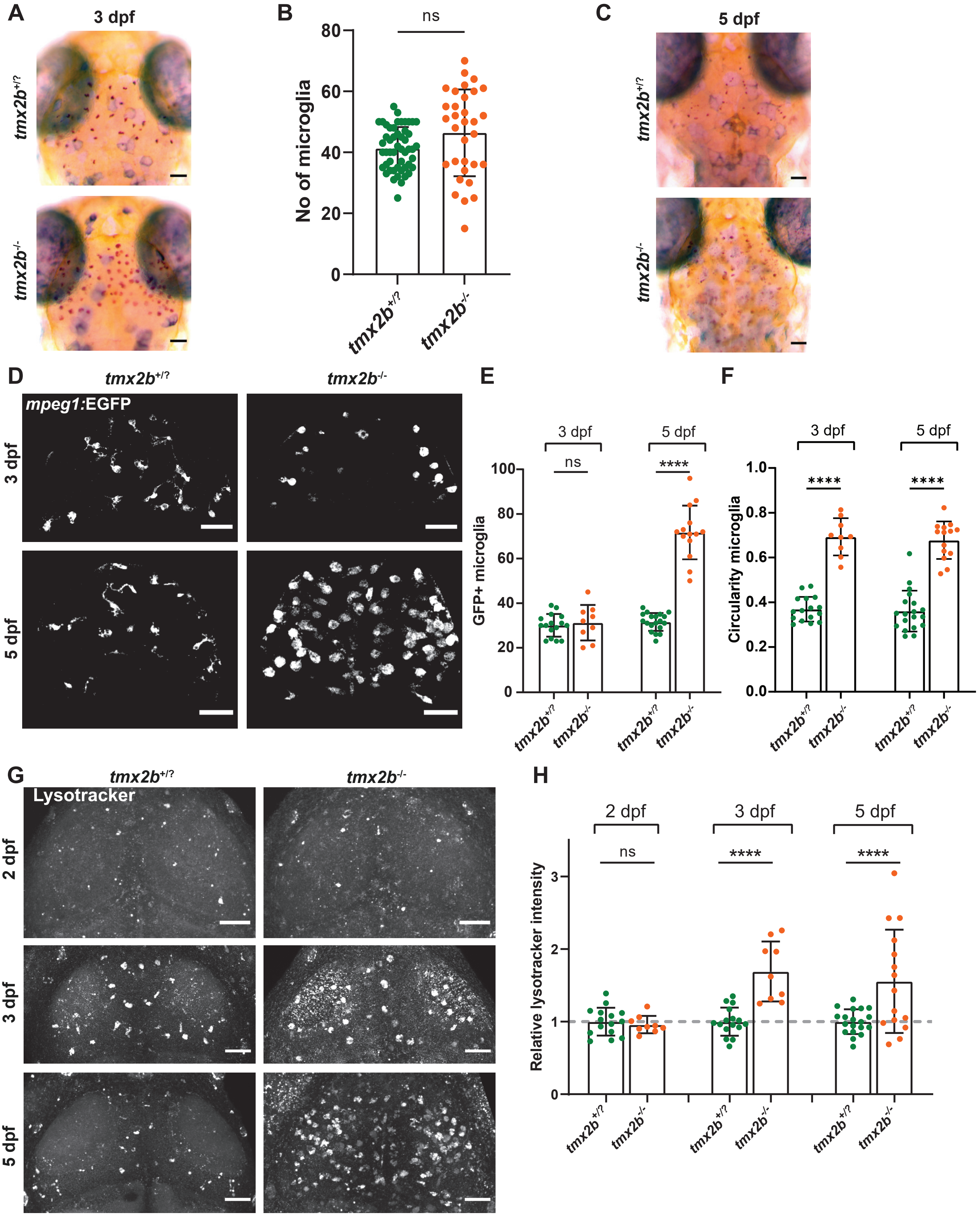


**Figure S11. Microglia activation and increased lysosome numbers in *tmx2b*$^{-/-}$ zebrafish (A)** Representative images of 3 dpf *tmx2b*$^{+/?}$ and *tmx2b*$^{-/-}$ zebrafish after neutral red (NR) staining. Scale bars indicate 100 µm. **(B)** Quantification of NR+ microglia at 3dpf. *tmx2b*$^{+/?}$ n=51 *tmx2b*$^{-/-}$ n=32. Unpaired t-test with Welch's correction. **(C)** Representative images of 5 dpf *tmx2b*$^{+/?}$ and *tmx2b*$^{-/-}$ zebrafish after neutral red (NR) staining. Scale bars indicate 100 µm. **(D)** Representative images of *mpeg1:*EGFP+ microglia in midbrain of *tmx2b*$^{+/?}$ and *tmx2b*$^{-/-}$ zebrafish at 3 and 5 dpf. Scale bars indicate 50 µm. **(E)** Quantification of *mpeg1:*EGFP+ microglia number in midbrain of t*mx2b*$^{+/?}$ and *tmx2b*$^{-/-}$ zebrafish at 3 and 5 dpf. **(F)** Circularity measurement of microglia in midbrain of *tmx2b*$^{+/?}$ and *tmx2b*$^{-/-}$ zebrafish at 3 and 5 dpf. Each dot represents the average value of six microglia from a single zebrafish brain. **(E,F)** 3,5 dpf: N=1,1 experiment, *tmx2b*$^{+/?}$ n=16,19, *tmx2b*$^{-/-}$ n=9,13 zebrafish. Two-way ANOVA, Šídák's multiple comparisons test. **(G)** Representative images of lysotracker+ lysosomes in midbrain of *tmx2b*$^{+/?}$ and *tmx2b*$^{-/-}$ zebrafish at 2,3 and 5 dpf. Scale bars indicate 50 µm. **(H)** Relative lysotracker fluoresence intensity normalized to the average *tmx2b*$^{+/?}$ at 2,3 or 5 dpf. 2,3,5 dpf: N=1.1,1 experiment, t*mx2b*$^{+/?}$ n=15,16,19, *tmx2b*$^{-/-}$ n=9,9,15 zebrafish. Two-way ANOVA, Šídák's multiple comparisons test. Data are represented as mean ± SD. *p < 0.05, **p < 0.01, ***p < 0.001, ****p < 0.0001.

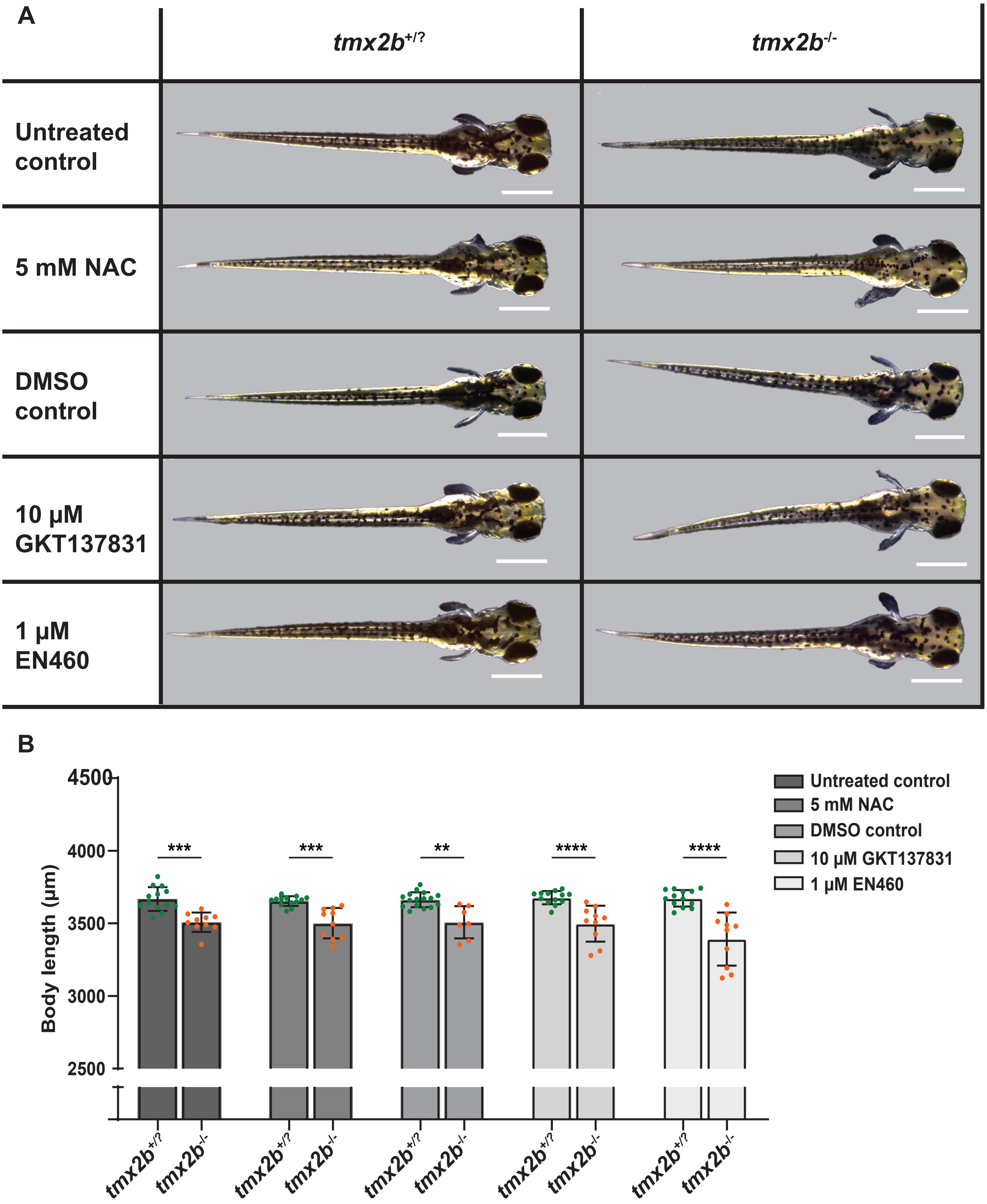


**Figure S12. ROS reducing drug treatments (A)** Representative images of 3 dpf *tmx2b*$^{+/?}$ and *tmx2b*$^{-/-}$ zebrafish treated with ROS reducing drugs. Scale bars indicate 500 µm. All treated *tmx2b*$^{-/-}$ zebrafish developed necrosis as is observed by an increased opacity in the brain compared to the *tmx2b*$^{+/?}$ zebrafish. **(B)** Body length measurements of 3 dpf zebrafish treated with ROS reducing drugs. Untreated, 5 mM NAC, DMSO control, 10 µM GKT137831, 1 µM EN460: N=1,1,1,1,1 experiment; *tmx2b*$^{+/?}$ , n=13,13,16,13,13 ; *tmx2b*$^{-/-}$, n=10,10,7,10,10 zebrafish. One-way ANOVA with Šídák's multiple comparisons test. Data are represented as mean ± SD. *$p < 0.05$, **$p < 0.01$, ***$p < 0.001$, ****$p < 0.0001$.

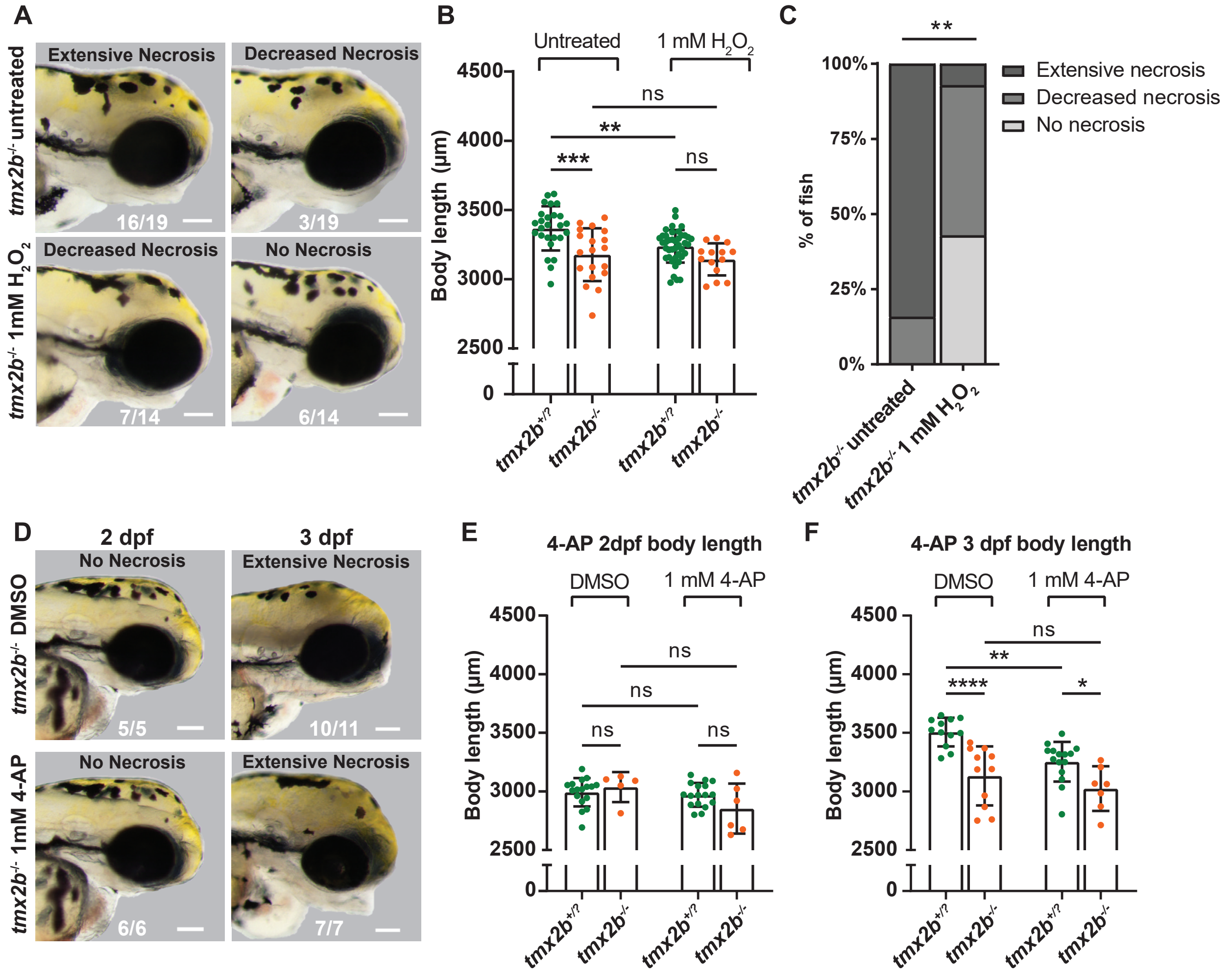


**Figure S13. $H_2O_2$ and 4-AP treatment. (A)** Brightfield images lateral view head of 3 dpf *tmx2b*$^{-/-}$ untreated (upper images) and treated with 1mM $H_2O_2$ (lower images). *tmx2b*$^{-/-}$ fish treated with 1mM $H_2O_2$ had either decreased necrosis (a reduced opacity compared to the extensive necrosis) or no necrosis at all. Bottom numbers indicate counts of zebrafish with specified genotype and phenotype. Scale bar represents 100 µm. **(B)** Body length measurements of 3 dpf untreated and 1mM $H_2O_2$ treated *tmx2b*$^{+/?}$ and *tmx2b*$^{-/-}$ zebrafish. Two-way ANOVA, Tukey's multiple comparisons test. **(C)** Quantification of **(A).** Fisher's exact test (extensive and decreased necrosis groups were combined for statistical test). **(B,C)** Untreated, 1mM H2O2 : N=2,2 experiments; *tmx2b*$^{+/?}$ , n=26,41 ; *tmx2b*$^{-/-}$ , n=19,14 zebrafish. **(D)** Brightfield images lateral view head of *tmx2b*$^{-/-}$ zebrafish untreated (upper images) and 1 mM 4-AP treated (lower images) at 2 and 3 dpf. Scale bar represents 100 µm. Bottom numbers indicate counts of zebrafish with specified genotype and phenotype. **(E)** Body length measurements of 2 dpf DMSO control and 1mM 4-AP treated zebrafish. DMSO control, 1mM 4-AP: N=1,1 experiment, t*mx2b*$^{+/?}$ n=17,17, t*mx2b*$^{-/-}$ n=5,6 zebrafish **(F)** Body length measurements of 3 dpf DMSO control and 1mM 4-AP treated zebrafish. DMSO control, 1mM 4-AP: N=1,1 experiment, t*mx2b*$^{+/?}$ n=12,15 t*mx2b*$^{-/-}$ n=11,7 zebrafish. Two-way ANOVA, Tukey's multiple comparisons test. Data are represented as mean ± SD. *$p < 0.05$, **$p < 0.01$, ***$p < 0.001$, ****$p < 0.0001$.

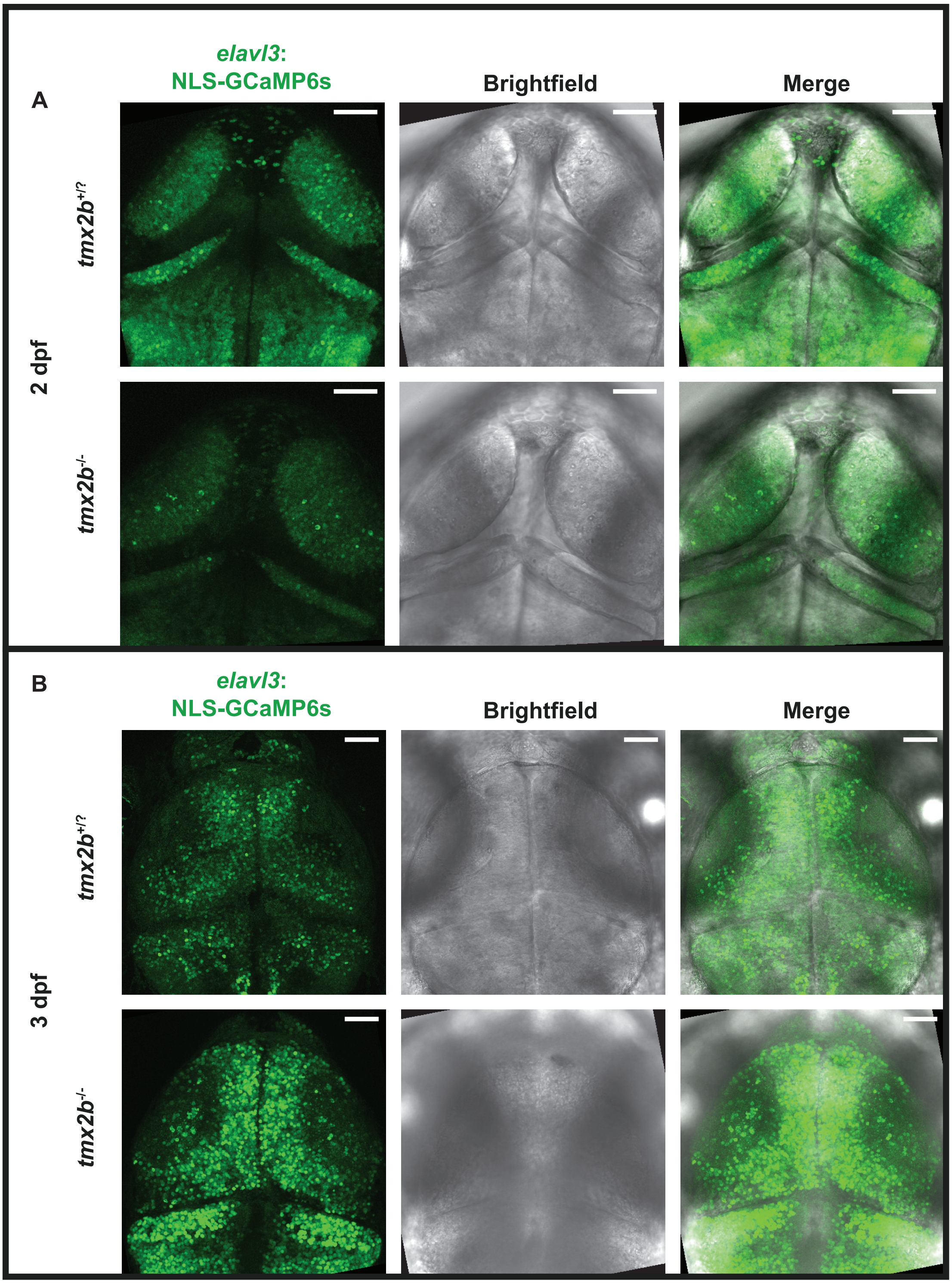


**Figure S14. Calcium imaging in *tmx2b*$^{-/-}$ zebrafish brain. (A,B)** Representative images of *elavl3:*NLS-GCaMP6s+ (green) neurons in t*mx2b*$^{+/?}$ and t*mx2b*$^{-/-}$ zebrafish at 2 and 3 dpf. 2 dpf: N=1 experiment, t*mx2b*$^{+/?}$ n=22, t*mx2b*$^{-/-}$ n=10 zebrafish. 3 dpf: N=1 experiment, t*mx2b*$^{+/?}$ n=16, t*mx2b*$^{-/-}$ n=16 zebrafish.
